# Pan-Cancer Analysis of DNA Methylation Identifies Genes and Biological Functions Associated with Overall Survival

**DOI:** 10.1101/2021.06.20.449136

**Authors:** Romola Cavet, Peng Yue, Guy Cavet

## Abstract

DNA methylation influences gene expression and is altered in many cancers, but the relationship between DNA methylation and cancer outcomes is not yet fully understood. If methylation of specific genes is associated with better or worse outcomes, it could implicate genes in driving cancer and suggest therapeutic strategies. To advance our understanding of DNA methylation in cancer biology, we conducted a pan-cancer analysis of the relationship between methylation and overall survival. Using data on 28 tumor types from The Cancer Genome Atlas (TCGA), we identified genes and genomic regions whose methylation was recurrently associated with survival across multiple cancer types. While global DNA methylation levels are associated with outcome in some cancers, we found that the gene-specific associations were largely independent of these global effects. Genes with recurrent associations across cancer types were enriched for certain biological functions, such as immunity and cell-cell adhesion. While these recurrently associated genes were found throughout the genome, they were enriched in certain genomic regions, which may further implicate certain gene families and gene clusters in affecting survival. By finding common features across cancer types, our results link DNA methylation to patient outcomes, identify biological mechanisms that could explain survival differences, and support the potential value of treatments that modulate the methylation of tumor DNA.

## Introduction

Cancer is defined by a plethora of genetic and epigenetic alterations that alter gene activity and promote uncontrolled cell proliferation [1]. As a consequence, the study of tumor genomes and large-scale computational data analysis have proved invaluable in understanding cancer development, prognosis, and treatment [2, 3].

DNA methylation is the epigenetic conversion of cytosine to 5-methylcytosine, and is widely implicated in both normal and abnormal regulation of genome function [4]. DNA methyltransferases (DNMTs) facilitate methylation, acting as either *de novo* DNMTs (which establish early methylation patterns during development) or as maintenance DNMTs (which copy patterns of DNA methylation onto new strands after DNA replication) [4, 5]. The mechanism by which DNMTs are regulated is suggested to be through their interaction with proliferating cell nuclear antigens (PCNA) and RAD9 [6].

DNA methylation has long been suspected to influence gene expression. Although the exact mechanism is yet unknown, one possibility is that it impedes transcription activators from binding to the DNA molecule [5]. DNA methylation of gene bodies and gene promoters have both been found to have an inverse relationship with gene expression [7, 8].

DNA methylation is frequently altered in cancer and is related both to tumorigenesis and prognosis. Both localized hypermethylation and global hypomethylation are observed in cancer and can be associated with the development and progression of the disease [9]. Hypermethylation in certain genomic regions can promote tumorigenesis [10]. For example, DNA methylation of promoter regions for tumor suppressor genes has repeatedly been implicated as an early driving factor in tumorigenesis [11–13]. DNA methylation inhibitors such as azacytidine and decitabine are proven treatments in some indications, and are believed to act by relieving inhibition of tumor suppressor gene transcription [14]. By contrast, global hypomethylation is associated with genomic instability and worse prognosis and/or progression in cancers including lower-grade glioma, prostate cancer, and breast adenocarcinoma [15–17].

Comparison of DNA methylation between tumor and normal tissue has identified patterns of aberrant methylation associated with cancer. For example, in a pan-cancer analysis of DNA methylation, patterns of DNA methylation instability were found in pathways related to cell metabolism, apoptosis, and DNA repair, such as CDKN1C, BRCA1, and MLH1 [18]. A pattern of hypermethylated genes encoding proteins in the WNT signaling pathway was also found in patients with various adenocarcinomas [19]. Identification of tumor DNA methylation patterns increases our disease knowledge and may facilitate early diagnosis [20, 21]. However, further analysis is required to determine whether and how these patterns affect cancer biology and disease outcomes.

We set out to directly seek methylation patterns associated with patient survival, with several goals. First, to assess the extent to which locus-specific methylation is linked to cancer outcomes. Second, to identify which genes have methylation consistently associated with survival across multiple tumor types. Finally, to identify higher-level biological functions and pathways that are enriched for these genes, and that have the potential to explain the relationships between methylation and clinical outcome. Our approach used large, publicly available cancer genomics datasets.

The Cancer Genome Atlas (TCGA) program makes available cancer genomic, epigenomic and clinical data available for over 20,000 samples spanning 33 different cancer types (https://www.cancer.gov/tcga). Using TCGA data we analyzed DNA methylation patterns measured in 28 different tumor types. This analysis identified genes whose methylation is associated with prognosis, both positively and negatively. Some of these genes have consistent associations across several types of cancer. These genes suggest biological mechanisms, shared across multiple tumor types, by which methylation may affect cancer outcomes.

## Materials and Methods

### Accessing Data

TCGA data were downloaded from https://www.cbioportal.org/. Firehose Legacy data sets were obtained for 32 cancer types, of which 4 were discarded due to missing data. Firehose Legacy data sets were chosen because they include large numbers of samples.

### Identifying genes for which methylation is associated with survival

Statistical analysis was conducted in the R programming language using the packages “survival,” “survminer,” and “dplyr”. Within each cancer type, the methylation hm450 data and patient clinical data were used to test for associations between gene-level methylation and overall survival. In the first analysis, a Cox proportional hazards model was fit to evaluate the association between each gene’s methylation level and overall survival. False discovery rates (*q*-values) were estimated using the method of Benjamini and Hochberg [22]. *Q*-values below 0.05 were considered significant. For each cancer type, the genes positively associated with survival (greater methylation associated with greater survival) and the genes negatively associated with survival (greater methylation associated with shorter survival) were considered separately in further analysis.

To evaluate associations between global methylation and survival for each cancer type, a global methylation score was calculated for each patient by taking the mean methylation score across all genes. Cox proportional hazards regression was used, for each cancer type separately, to evaluate the association between global methylation score and overall survival.

To determine whether individual genes’ methylation levels were still associated with survival when taking into account global methylation, another Cox proportional hazards regression analysis was carried out for each gene that had a *q*-value < 0.05 in the first analysis. For each cancer type and for each gene, the regression evaluated the association with overall survival, now including both the individual gene’s methylation values and the global methylation scores. *P*-values < 0.05 for the individual gene’s methylation were considered significant.

### Finding the association between methylation and mRNA expression

For each study with genes whose methylation was significantly associated with survival, the relationships between RNA-Seq values (all sample median Z scores) and methylation were evaluated by Spearman correlation.

### Gene set enrichment analysis

The MSigDB annotate tool (http://www.gsea-msigdb.org/gsea/msigdb/annotate.jsp) [23, 24] was used to evaluate overlap between genes recurrently associated with survival (*q* < 0.05 in 3 or more studies) and gene sets based on the Gene Ontology, “C5 GO: Gene Ontology” [25, 26]. Genes positively associated with survival (in ≥ 3 studies) and genes negatively associated with survival (in ≥ 3 studies) were analyzed separately.

### Determining enriched chromosomal locations

Gene chromosomal locations and chromosome lengths were obtained from the Genome Reference Consortium (https://www.ncbi.nlm.nih.gov/grc/human/data). Each chromosome was scanned using a window size of 5 × 10^6^ bases, shifted along the chromosome 1 × 10^5^ bases at a time. For each window position, the number of recurrent genes within the window was calculated and used to identify regions with the greatest density.

## Results

### Identification of genes whose methylation is associated with survival

In order to identify genes and mechanisms linking DNA methylation to cancer outcome, we first identified genes for which methylation was significantly associated with overall survival for each of 28 TCGA studies representing different cancers. The numbers of genes with such associations varied widely between tumor types, from 0 to 8,781, with a median of 124.5 (Table 1, S1 Table, S2 Table). This variation between studies is likely influenced by differences between cancer types in the biological role of methylation, but also differences in sample size and statistical power.

**Table 1.** Associations between gene-level methylation and survival by tumor type. Numbers of genes represent the total number whose methylation was associated with survival (either greater or shorter).

| Tumor type | Abbreviation | Number of events/patients | Number of genes with associations | Associations when considering global methylation | Association between global methylation and survival (*p*-value) |
| --- | --- | --- | --- | --- | --- |
| Brain lower grade glioma | LGG | 29/80 | 8781 | 3701 | $1.9 \times 10^{-20}$ |
| Kidney renal clear cell carcinoma | KIRC | 182/412 | 4394 | 4373 | 0.089 |
| Kidney renal papillary cell carcinoma | KIRP | 104/783 | 4157 | 3980 | 0.049 |
| Skin cutaneous melanoma | SKCM | 72/307 | 1286 | 1286 | 0.35 |
| Liver hepatocellular carcinoma | LIHC | 18/36 | 1259 | 1259 | 0.096 |
| Uveal melanoma | UVM | 88/390 | 1087 | 1087 | 0.78 |
| Adrenocortical carcinoma | ACC | 9/48 | 1086 | 1013 | 0.015 |
| Uterine corpus endometrial carcinoma | UCEC | 77/185 | 663 | 663 | 0.90 |
| Cervical squamous cell carcinoma and endocervical adenocarcinoma | CESC | 94/139 | 644 | 644 | 0.66 |
| Thymoma | THYM | 224/528 | 493 | 492 | 0.042 |
| Pancreatic adenocarcinoma | PAAD | 10/66 | 454 | 454 | 0.99 |
| Kidney chromophobe | KICH | 105/319 | 258 | 258 | 0.75 |
| Lung adenocarcinoma | LUAD | 40/275 | 178 | 178 | 0.37 |
| Sarcoma | SARC | 129/194 | 130 | 130 | 0.78 |
| Thyroid carcinoma | THCA | 126/515 | 119 | 117 | 0.016 |
| Breast invasive carcinoma | BRCA | 132/377 | 110 | 110 | 0.27 |
| Bladder urothelial carcinoma | BLCA | 165/458 | 97 | 97 | 0.69 |
| Head and neck squamous cell carcinoma | HNSC | 160/370 | 80 | 80 | 0.31 |
| Acute myeloid leukemia | LAML | 99/184 | 3 | 3 | 0.25 |
| Esophageal carcinoma | ESCA | 10/498 | 2 | 2 | 0.44 |
| Colon adenocarcinoma | COAD | 99/261 | 1 | 1 | 0.078 |
| Cholangiocarcinoma | CHOL | 223/470 | 0 | 0 | 0.17 |
| Lung squamous cell carcinoma | LUSC | 155/395 | 0 | 0 | 0.86 |
| Testicular germ cell tumors | TGCT | 4/134 | 0 | 0 | 0.63 |
| Prostate adenocarcinoma | PRAD | 16/503 | 0 | 0 | 0.45 |
| Lymphoid neoplasm diffuse large b-cell lymphoma | DLBC | 9/124 | 0 | 0 | 0.092 |
| Stomach adenocarcinoma | STAD | 73/431 | 0 | 0 | 0.083 |
| Glioblastoma multiforme | GBM | 23/80 | 0 | 0 | 0.0098 |

### Most associations with survival are gene- or region-specific

The goal of this study was to seek genes whose methylation was related to patient outcome due to the specific biology of those genes (for example, genes that promote or inhibit metastasis). However, global methylation level is a prognostic factor in certain cancer types, which could result in widespread gene-level associations without regard to the biology of the individual genes. For example, global hypermethylation is observed in a subset of gliomas that have relatively favorable prognosis, often linked to IDH1 mutation [27]. To take this into account, we first evaluated whether the average genome-wide methylation level was associated with survival for each cancer type. We observed significant associations (p<0.05) for 6 cancer types, including lower-grade glioma and glioblastoma (Table 1). This may help explain the very large number of genes whose methylation was associated with survival in lower-grade glioma (8,781), and it could also be a contributor to the results observed in a few other cancer types (such as renal clear cell and papillary cell carcinomas). To ensure that our gene-specific associations were not being dominated by the effect of global methylation, we analyzed whether each gene’s methylation was still associated with survival when global methylation was taken into account. For lower-grade glioma, many genes were no longer significantly associated with survival, but for all other cancer types, the vast majority of gene-level associations were still significant, even after taking global methylation into account (Table 1). This suggests that these associations with outcome were due to gene-specific or region-specific mechanisms.

### Gene-level methylation is associated with lower mRNA levels

Methylation has long been associated with transcriptional silencing [4]. We confirmed that methylation of individual genes in the data sets we used was overwhelmingly negatively correlated with mRNA levels (Fig 1). The distribution of correlation coefficients was very consistent across tumor types (S1 Fig). This negative correlation is consistent with gene-level methylation influencing patient outcomes by reducing transcription.

**Fig 1.**
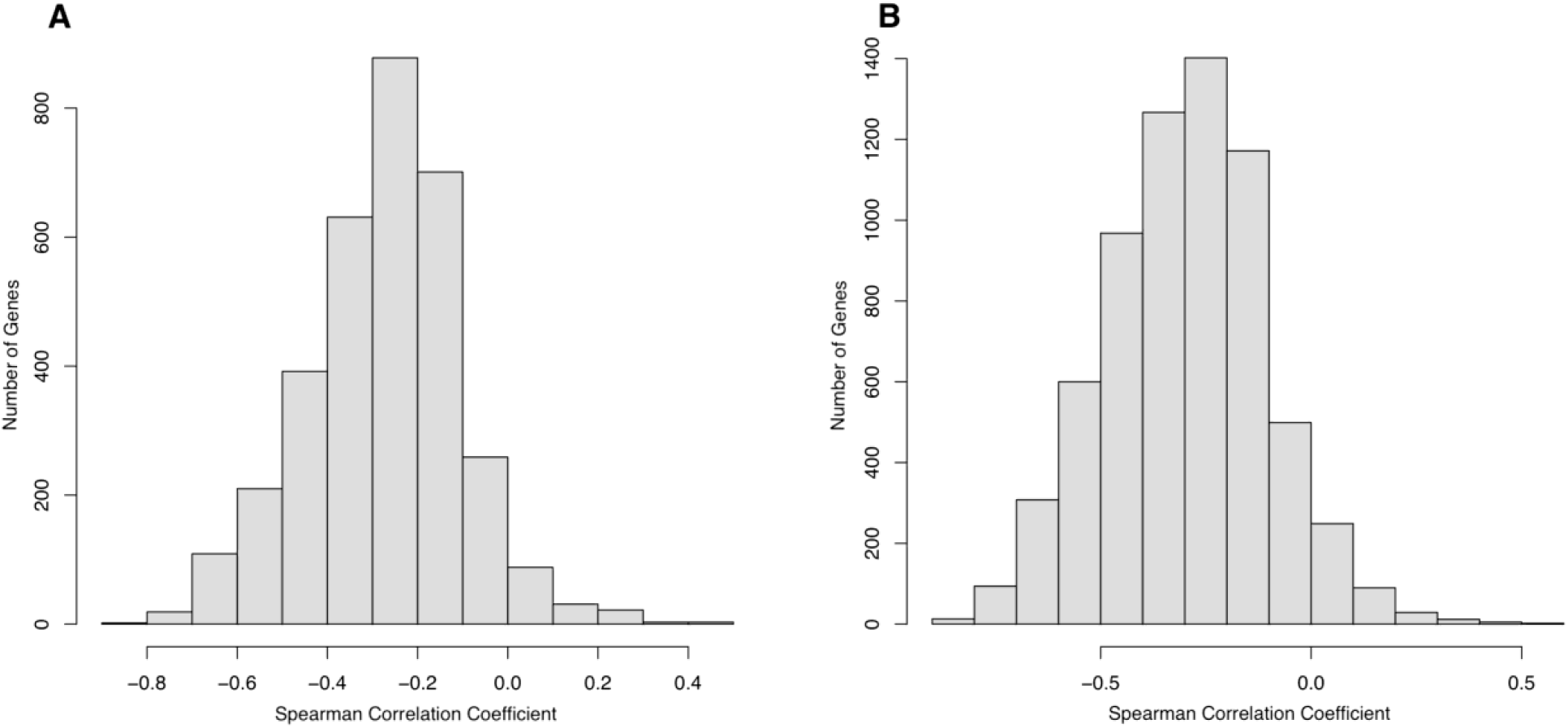
Most genes’ expression is inversely correlated with their methylation. Representative histograms of Spearman correlation between gene methylation and RNA-Seq expression values for (A) kidney renal clear cell carcinoma and (B) lower grade glioma.

### Chromosomal regions enriched in survival-associated genes

The identification of genes whose methylation is associated with survival raises the question: are these genes clustered in certain genomic regions that are methylated in unison? We found that genes whose methylation was recurrently associated with survival (across 3 or more cancer types) were scattered throughout the genome, but there were certain regions in which they occurred with greater density. While some variation in density will occur by chance, these regions could reflect biological mechanisms connecting methylation to cancer biology.

When considering genes whose methylation was associated with shorter survival, a prominent density peak was identified around 33Mbp on chromosome 6 (Fig 2A). This peak coincides with the HLA locus, which encodes many receptors critical to the adaptive immune system. This peak had the second-highest density of negatively-associated genes across the entire genome. The ten recurrently associated genes within this region were HLA-E, HLA-B, LST1, TNXB, HLA-DRA, HLA-DQA1, PSMB9, HLA-DMA, HLA-DPB2, and HMGA1. These include genes encoding MHC class I receptors, class II receptors, and proteins with other immune-related functions. The region with the highest density genome-wide was centered around 156Mbp on chromosome 1q (Fig 2B). The functions of the genes in this region were more heterogeneous than those under the peak on chromosome 6, but were also enriched in genes implicated in adaptive immunity including CD1c, CD244, SLAMF7, and LY9/SLAMF3 (S3 Table).

**Figure 2.**
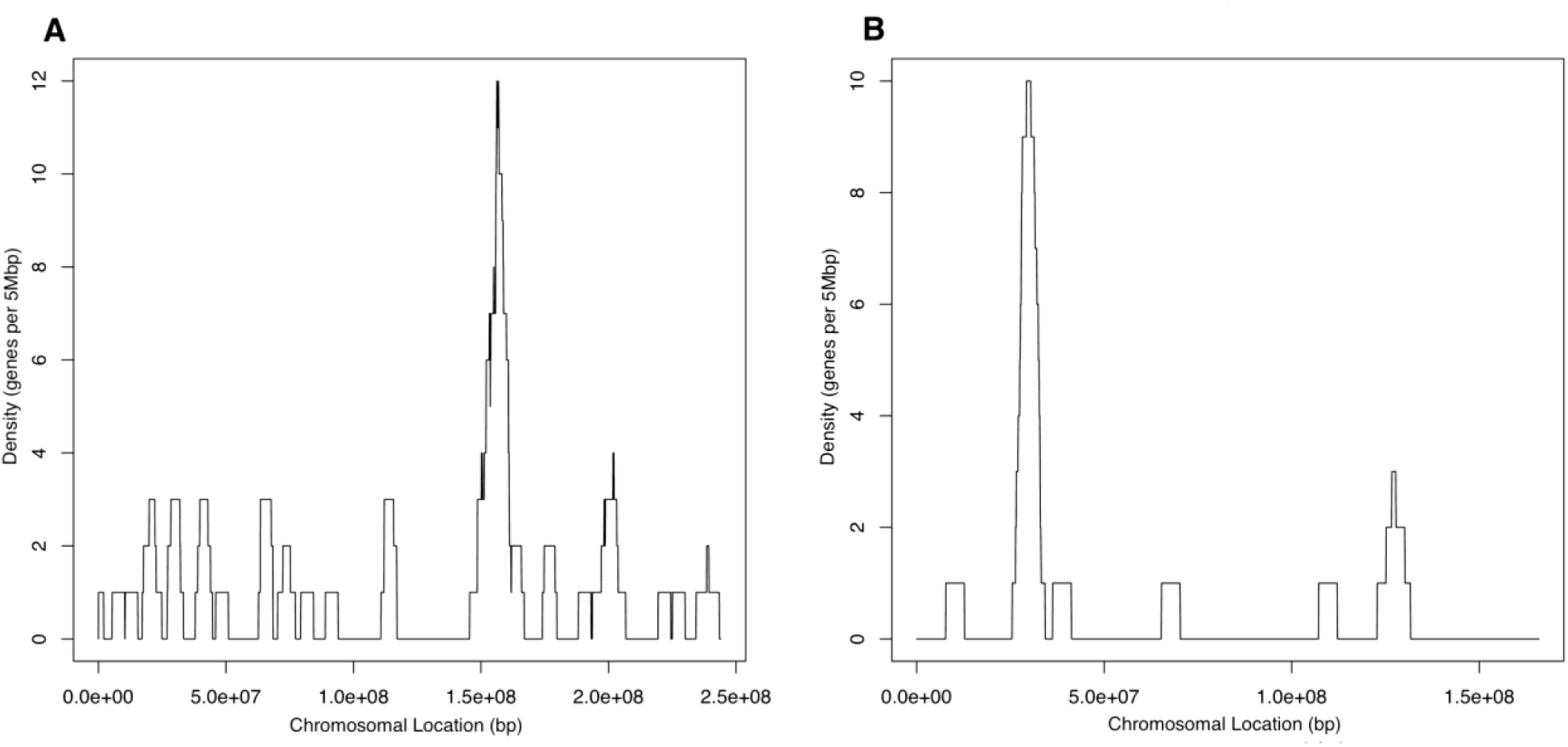
Chromosomes with highest density of genes with methylation recurrently associated with shorter survival (q < 0.05 in at least 3 cancers). Plots indicate number of genes with associations within a 5 Mbp window on (A) chromosome 6, (B) chromosome 1.

In the analysis of genes whose methylation was positively associated with survival, the genomic region with the highest density was around 57Mbp on chromosome 19q (Fig 3A). There were 14 recurrent genes within this region, all of which encode zinc finger motifs: ZNF582, ZFP28, ZNF835, ZNF304, ZNF549, ZIK1, ZNF154, ZNF671, ZSCAN1, ZNF135, ZSCAN18, ZNF274, ZNF544, and ZNF132. The region represents one of several clusters of zinc finger genes on chromosome 19, many of which encode transcription factors that have been linked to the development or progression of cancer [28, 29]. The region with the second highest density of genes with positive associations was centered around 6Mbp on chromosome 17p (Fig 3B). The 8 genes identified in this region include 2 that also encode zinc finger proteins (ZFP3, ZBTB4) but otherwise have more diverse functions (S3 Table). Further analysis of these genes may identify other potential mechanisms linking methylation to outcome.

**Fig 3.**
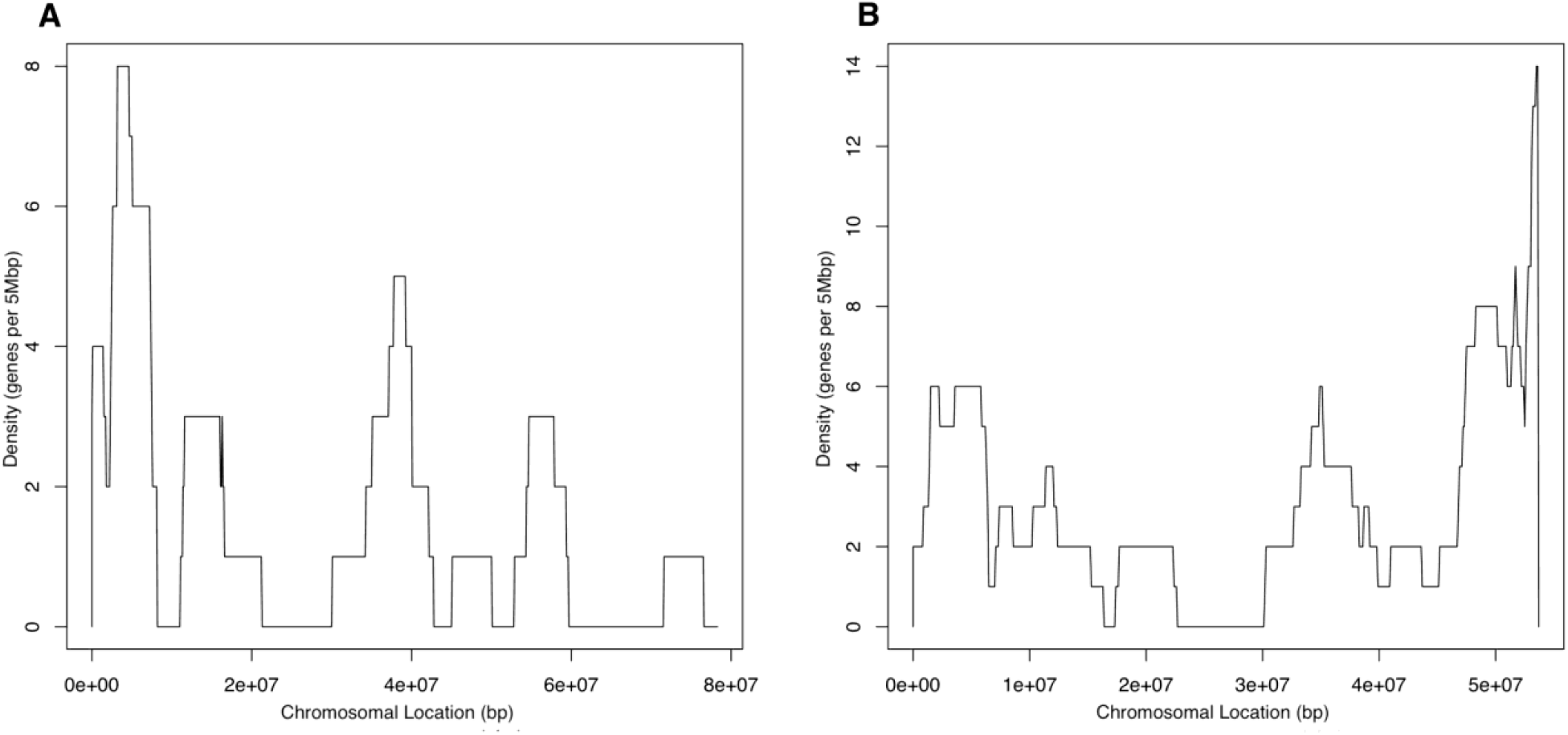
Chromosomes with highest density of genes with methylation recurrently associated with greater survival (q < 0.05 in at least 3 cancers). Plots indicate number of genes with associations within a 5 Mbp window on (A) chromosome 19, (B) chromosome 17.

### Survival-associated genes are enriched for the immune system, cell activation, and adhesion

To identify pathways and gene functions that could explain connections between methylation and survival, we identified Gene Ontology gene sets that were over-represented among the genes whose methylation was associated with survival across cancer types. Genes whose methylation was negatively associated with survival were strongly enriched in immune system genes sets (Table 2, S4 Table). Of the ten gene sets that had the most significant overlaps, four were involved in immune system regulation, activation, and signaling. There were 96 genes overlapping with at least one immune-related gene set, including 6 in the HLA locus, several interleukin signaling genes (IL1A, IL10, IL6, IL20RB, IL2RA), and other immune functions. Apart from immune-related gene sets, the most significantly enriched biological processes were Cell Activation (signaling-related genes, relevant to activation of immune cells and also of other cell types) and Biological Adhesion (including classic adhesion molecule genes such as cadherins and integrins, and also genes involved in control of adhesion such as ADAM12, ADAM8 and ADAMDEC1). A comparison across tumor types revealed which tumor types were most enriched for each of the top gene sets (Fig 4). For example, genes identified in uveal melanoma were particularly enriched for immune-related gene sets, suggesting methylation of immune-related genes may be strongly associated with poor outcomes in patients with this cancer.

**Fig 4.**
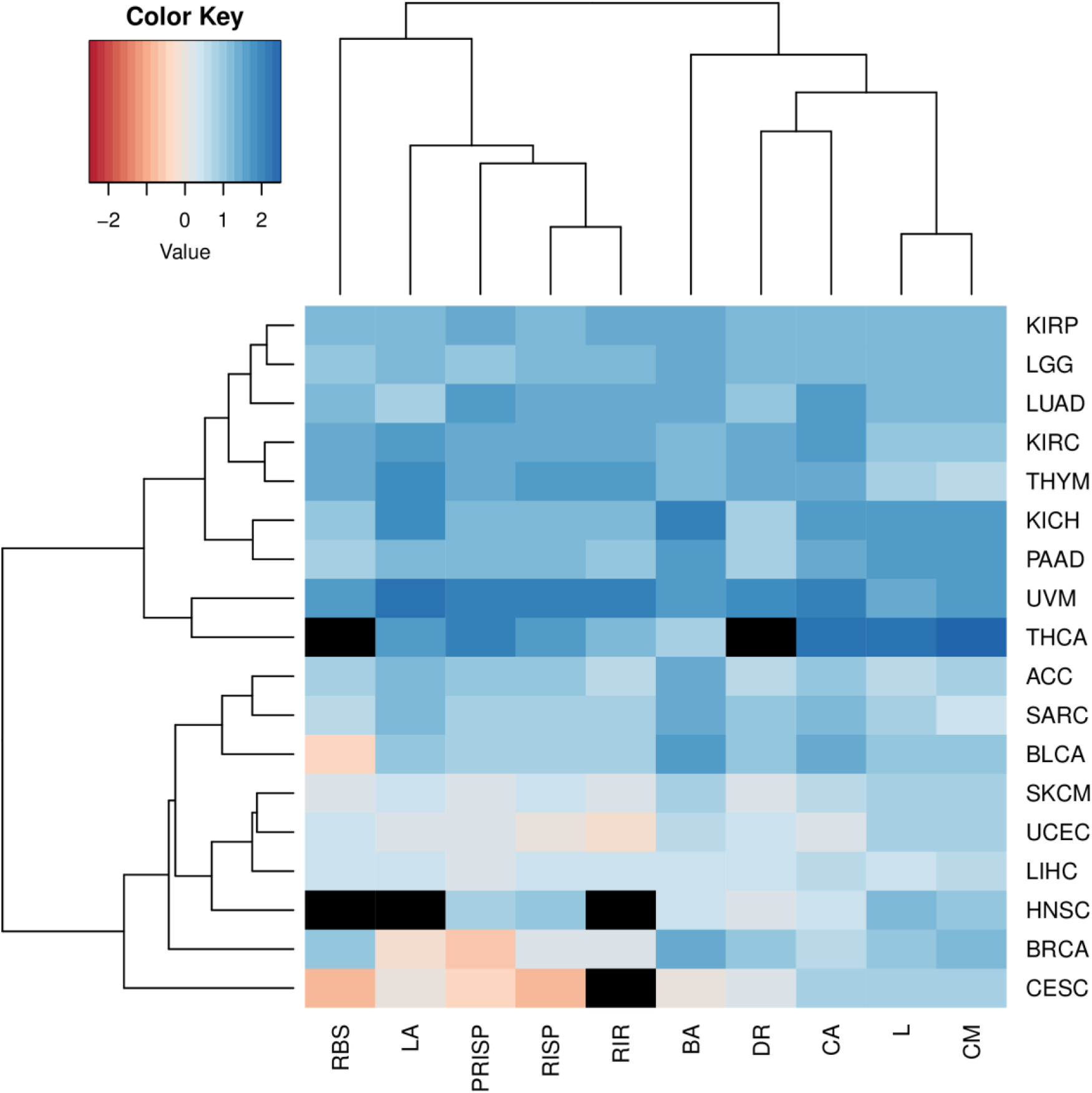
Gene set enrichments by cancer type, for genes associated with shorter survival. Columns represent GO gene sets; rows represent tumor types. Color indicates degree of enrichment (natural log odds ratio for the overlap between negatively associated genes for each study and each of ten GO gene sets). See Table 2 for gene set abbreviations. Blue indicates genes identified in a tumor type are enriched for a gene set, and red indicates that they are depleted. Black represents N/A.

**Table 2.** The ten gene sets most significantly enriched among genes whose methylation is associated with shorter survival.

| Gene Set Name | GO Identifier | Abbreviation | Number of Genes in Overlap | *Q*-value |
| --- | --- | --- | --- | --- |
| Cell Activation | GO:0001775 | CA | 83 | $5.5 \text{ e}^{-31}$ |
| Regulation of Immune System Process | GO:0002682 | RISP | 86 | $6.05 \text{ e}^{-31}$ |
| Lymphocyte Activation | GO:0046649 | LA | 54 | $2.74 \text{ e}^{-24}$ |
| Biological Adhesion | GO:0022610 | BA | 73 | $1.38 \text{ e}^{-23}$ |
| Cell Migration | GO:0016477 | CM | 76 | $1.38 \text{ e}^{-23}$ |
| Locomotion | GO:0040011 | L | 84 | $2.26 \text{ e}^{-23}$ |
| Regulation of Immune Response | GO:0050776 | RIR | 58 | $7.12 \text{ e}^{-22}$ |
| Positive Regulation of Immune System Process | GO:0002684 | PRISP | 59 | $1.67 \text{ e}^{-21}$ |
| Response to Biotic Stimulus | GO:0009607 | RBS | 71 | $2.36 \text{ e}^{-20}$ |
| Defense Response | GO:0006952 | DR | 74 | $1.15 \text{ e}^{-19}$ |

Genes whose methylation was associated with greater survival were strongly enriched for transcription factors (TFs) and other regulators of mRNA transcription (Table 3, S5 Table). The two GO gene sets with most statistically significant enrichment were Transcription Regulatory Activity and DNA Binding Transcription Factor Activity, and many other highly ranked gene sets were also transcription-related. The overlapping genes included many encoding zinc fingers, such as the cluster on chromosome 19 mentioned above, but also other TF families such as helix-loop-helix, forkhead, and homeobox (S5 Table). While individual TFs can have both oncogenic and tumor suppressor activity, this result suggests that greater methylation of many TF genes is a favorable prognostic marker. Comparing across tumor types revealed which tumor types were most enriched for each of the top gene sets (Fig 5). For example, genes identified in breast cancer were particularly enriched for many of the sets, suggesting methylation of genes involved in transcriptional regulation may be strongly associated with better outcomes in breast cancer patients.

**Figure 5.**
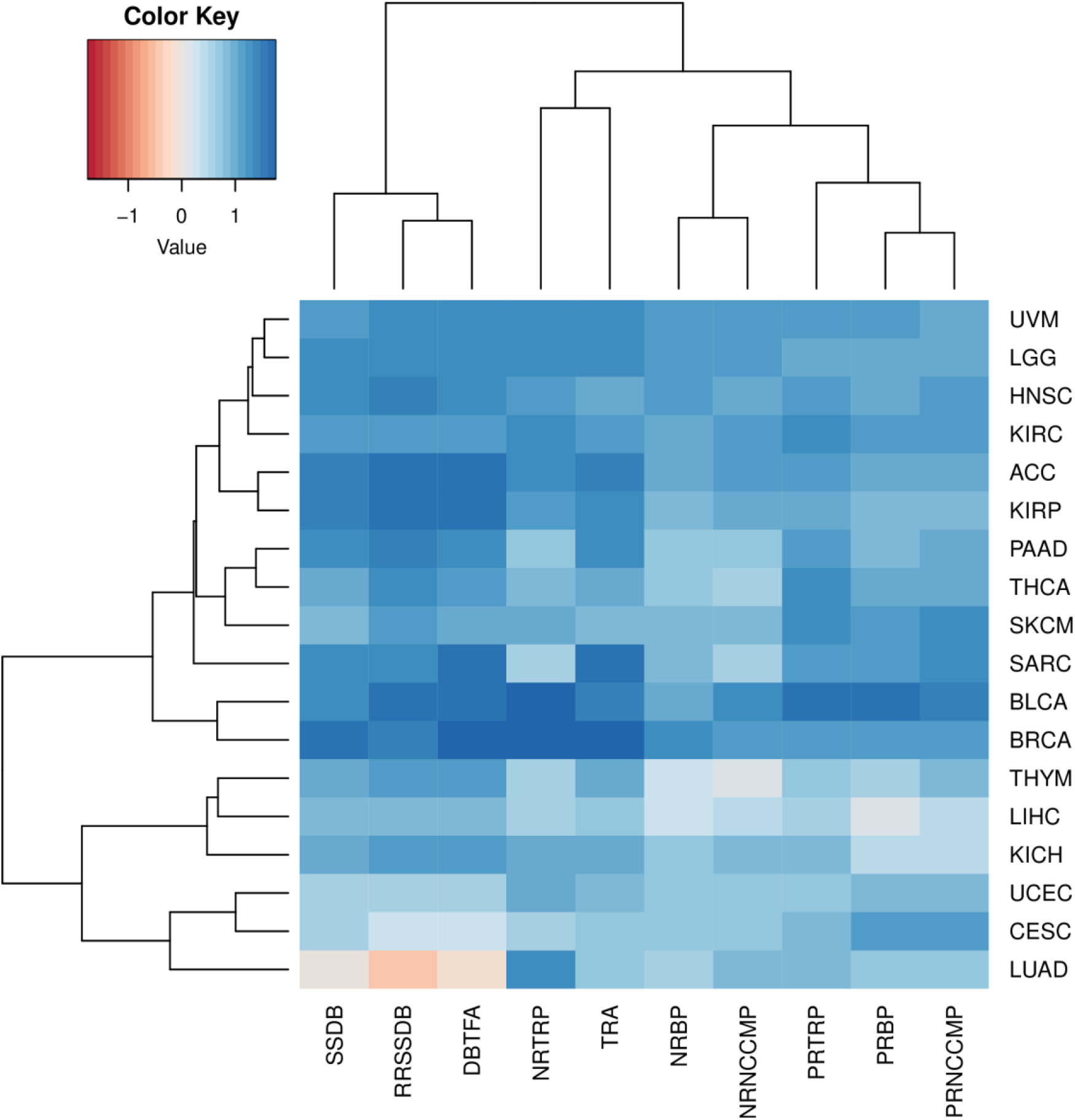
Gene set enrichments by cancer type, for genes associated with greater survival. Columns represent GO gene sets; rows represent tumor types. Color indicates degree of enrichment (natural log odds ratio for the overlap between negatively associated genes for each study and each of ten GO gene sets). See Table 2 for gene set abbreviations. Blue indicates genes identified in a tumor type are enriched for a gene set, and red indicates that they are depleted.

**Table 3.** The ten gene sets most significantly enriched among genes whose methylation is associated with greater survival.

| Gene Set Name | GO Identifier | Abbreviation | Number of Genes in Overlap | FDR q-value |
| --- | --- | --- | --- | --- |
| Transcription Regulator Activity | GO:0140110 | TRA | 90 | $1.85 \times 10^{-30}$ |
| DNA Binding Transcription Factor Activity | GO:0003700 | DBTFA | 78 | $2.94 \times 10^{-30}$ |
| Sequence Specific DNA Binding | GO:0043565 | SSDB | 82 | $5.86 \times 10^{-29}$ |
| Cis Regulatory Region Sequence Specific DNA Binding | GO:0000987 | RRSSDB | 70 | $2.18 \times 10^{-28}$ |
| Negative Regulation of Nucleobase Containing Compound Metabolic Process | GO:0045934 | NRNCCMP | 66 | $7.14 \times 10^{-20}$ |
| Positive Regulation of Nucleobase Containing Compound Metabolic Process | GO:0045935 | PRNCCMP | 70 | $6.9 \times 10^{-18}$ |
| Negative Regulation of Biosynthetic Process | GO:0009890 | NRBP | 66 | $1.31 \times 10^{-17}$ |
| Negative Regulation of Transcription by RNA Polymerase II | GO:0000122 | NRTRP | 47 | $3.06 \times 10^{-17}$ |
| Positive Regulation of Biosynthetic Process | GO:0009891 | PRBP | 69 | $3.29 \times 10^{-16}$ |
| Positive Regulation of Transcription by RNA Polymerase II | GO:0045944 | PRTRP | 50 | $3.11 \times 10^{-14}$ |

## Discussion

DNA methylation has long been known to be dysregulated in cancer, but its relationship to patient outcomes is only partly understood [30]. The differences in methylation pattern between tumor and normal tissue have been described in detail and may provide clues to the origins of the disease [30]. However, studying the relationship between methylation and cancer outcomes requires a different approach and addresses a different question: among patients who have cancer of a given type, what genes and biological processes connect methylation to outcome? We conducted a pan-cancer analysis to identify genes whose methylation is associated with overall survival, and analyzed potential biological explanations. This approach found genes whose methylation is consistently associated with survival across multiple cancer types, and highlighted potential mechanisms.

Gene-specific methylation in tumors may affect patient outcomes by suppressing transcription and thereby changing tumor biology [5, 30]. In accordance with expectations, we found widespread inverse correlation between gene methylation and mRNA levels, consistent with the model that methylation can affect survival by reducing gene expression. Under this model, methylation of tumor suppressor genes would be associated with shorter survival, and methylation of oncogenes would be associated with greater survival. However, we cannot assume that an association between a gene’s methylation and survival means that gene is a “driver” that causes better or worse patient outcomes. There are other potential explanations for such associations. One of these is a relationship between outcome and the global level of genome methylation, as is observed in low grade glioma (LGG). We found that the known relationship between genome-wide DNA methylation and survival in LGG could account for many of the gene-level associations in that tumor type. However, LGG was exceptional in this regard: almost all of the gene-level associations we identified in other tumor types remained significant even when global methylation n was taken into account, suggesting those associations reflect by gene- or region-specific methylation. This does not mean every gene whose methylation is associated with survival is playing a causal role in determining patient outcomes. Such associations could also arise through confounding, for example if methylation and survival were being influenced by a third factor that affects both (such as a mechanism driving genomic instability). In addition, some genes could exhibit significant associations as a result of being located close to “driver” genes and having correlated methylation levels. This may be the case, for example, in the HLA locus. Within this locus our analysis highlighted class I MHC genes, hypermethylation of which has previously been reported to impair anti-tumor immune responses in multiple cancer types by reducing presentation of tumor-specific peptides [31]. We also found some nearby class II MHC genes had methylation associated with shorter survival. These may be “passenger” genes, in the sense that they that have methylation levels correlated to those of their class I neighbors, but no role in tumor immunity. On the other hand, class II MHC on tumors has also been reported to contribute to effective anti-tumor immune responses [32], so the methylation patterns we identified in these class IKI genes may also be functionally important. Other nearby genes with significant associations include PSMB9, which is involved in immunoproteasome digestion of peptides for presentation on MHC class I [33, 34] and LST1, which is implicated in immunity to pathogens and self antigens, and which may facilitate transfer of class I MHC molecules [35, 36]. The connections between HLA locus methylation and patient outcomes remain to be fully defined, but may represent additional opportunities for immunotherapy through modulation of methylation.

While significant associations were observed between gene methylation and outcomes in most cancer types, there were six cancer types in which no such associations were found: cholangiocarcinoma, diffuse large b-cell lymphoma, glioblastoma multiforme, lung squamous cell carcinoma, stomach adenocarcinoma, and testicular germ cell tumor. In some cases this is likely due, at least in part, to lack of statistical power. The cholangiocarcinoma and diffuse large b-cell lymphoma datasets had only 36 and 48 patients, respectively. In the case of testicular germ cell tumors, there were only 4 events. However, lack of statistical power is less likely to explain the lack of gene-specific associations in glioblastoma multiforme, lung squamous cell carcinoma, and stomach adenocarcinoma, which had larger numbers of samples and events. It may be that methylation of specific loci does not play such an important role in the biology of these cancers. The results for lung squamous cell carcinoma (LUSC) contrasted with those for lung adenocarcinoma, in which 178 genes were found to have associations between methylation and survival, despite being represented by fewer samples and fewer events than LUSC.

The contrast between glioblastoma multiforme and lower-grade glioma is also striking. Global methylation was significantly associated with survival in both cancers, as reported previously [16, 37]. However, there were no genes with significant associations in glioblastoma patients, while there were thousands with associations in lower-grade glioma patients, even after taking global methylation into account. This suggests there may be more potential to benefit patients by localized modulation of methylation in lower-grade gliomas than in glioblastoma.

Many genes for which DNA methylation was associated with greater survival were transcription factors or otherwise involved in regulation of transcription. Among these were 50 zinc finger genes. Zinc finger proteins are known to be involved in tumorigenesis, cancer progression, metastasis, and development/differentiation [38]. However, they act to suppress tumors in some circumstances, and to promote them in others. The large number of zinc finger genes in our results suggests that DNA methylation and suppression of these particular genes is beneficial. Twenty-two of these zinc finger genes were located on chr19q13, and may be affected by coordinated methylation of this region.

The genes whose methylation was associated with better outcomes also included 5 members of the forkhead box (FOX) family. The overactivity of FOX proteins is frequently associated with cancer progression [39], consistent with the idea that methylation of FOX genes could benefit patients. The FOX genes that recurred with positive associations between methylation and survival have almost all been shown to be implicated in cancer progression (the exception, FOXD4L6, has not been heavily studied). FOXC2 has been shown to be tumorigenic by promoting cell proliferation and carcinogenesis, and is overexpressed in breast, stomach, lung, prostate, cervical, and ovarian cancers [39, 40]. FOXE1’s effect on cancer is under debate, as it has been suggested to both inhibit proliferation and worsen prognosis [41, 42]. FOXF1 is associated with increased metastasis and progression in prostate cancer [39]. FOXQ1 has been shown to promote the progression of esophageal, breast, pancreatic, and colorectal cancers [39].

Genes with recurrent associations between methylation and positive outcome also included four members of the MAPK family, which mediate mitogenic signaling and have long been implicated in metastasis and cancer progression [43]. Three of these four MAPK genes have been shown to promote malignancy and cancer progression: MAP2K5 promotes epithelial cell malignant transformation, MAPK15 promotes cell proliferation in testicular germ cell tumors, and MAPK4 is tumorigenic in lung adenocarcinoma, bladder cancer, low-grade glioma, and thyroid carcinoma [44–47]. Interestingly, MAP2K3 has been shown to promote senescence [48].

Genes with recurrent associations between methylation and worse outcomes were most strongly enriched in immune system functions. These include class I MHC genes, which are known to be transcriptionally repressed by methylation in some tumor types, but also genes involved in other facets of the immune system, such as class II MHC, interleukin signaling (e.g., IL1A, IL10, IL6, IL1RAP, IL2RA), chemokine signaling (e.g., CCR2, CCR7, CCL3, CCRL2, CXCR6), and toll-like receptors (TLR1, TLR6). Each of these components of the immune system has its own complex connections to cancer biology. For example the interleukin 1 family can be both anti-tumorigenic and tumorigenic; it has been shown to promote inflammation and tumor growth in some cancers but the opposite has been shown in others [49]. Interleukin 2 (IL2) is a cytokine crucial in promoting T-cell and NK cell cytotoxicity and T-cell differentiation, and is an important currently used cancer immunotherapy due to its anti-tumor effects [50]. Interleukin 6 is an inflammatory cytokine whose overexpression has been shown to be detrimental to cancer patients’ outcomes [51]. Interleukin 10 is commonly thought of as an antiinflammatory cytokine, yet IL-10 deficiency has been linked to rapid tumor development and overexpression has been shown to lead to tumor rejection and tumor immunity in mice and humans [52]. The breadth and diversity of the connections between immune-related gene methylation and survival suggest that epigenetics may play a broader role in regulating anti-tumor immunity and determining outcomes than has hitherto been appreciated.

Genes whose methylation was recurrently associated with shorter survival were also strongly enriched for functions in cell-cell adhesion. Dysregulation of cell-cell adhesion is a hallmark of cancer, associated with the epithelial-mesenchymal transition, loss of anchorage-dependent growth, and metastasis [1, 53]. The genes whose methylation was associated with worse outcomes include those encoding adhesion molecules such as caherins (CDH3, CDH13, CDH20) and integrins (ITGAL, ITGB5, ITGAE), but also protein families with other roles in cancer biology such as ADAM metallopeptidases [54, 55] and desmocollin 2, whose downregulation is associated with metastasis [56, 57].

The approach used in this study is subject to a number of limitations. While TCGA offers extensive and rich data, it does not include all cancer types, and some of the studies had relatively few patients or events, which limited statistical power. In addition, our approach of analyzing gene-level methylation does not take into account potential differences in methylation between regions of a gene, such as gene bodies and promoters. Finally, while our results support the idea that local methylation is affecting patient outcomes by regulating specific biological processes, they do not establish causal links between methylation and outcome. Our findings provide a basis for further research in a number of directions, including more detailed examination of pathways and regions that are recurrently affected by methylation and associated with patient survival. In particular, our results establish evidence-based hypotheses for how methylation may affect outcome that may be tested rigorously with prospective laboratory and clinical studies.

Our results suggest DNA methylation may influence overall survival in more ways than were previously understood, and they pinpoint biological mechanisms that might mediate these effects. If further research confirms and extends our findings, the resulting knowledge may inform improved treatments aimed at improving patient outcomes. Some of these treatments may act by modulating methylation, including existing drugs such as azacitidine and decitabine, but greater promise may lie in approaches that alter methylation in a locus-specific manner. Sequence-specific methylation modulators are under development, for example by engineering methyltransferases with Cas9 protein components [58]. These technologies may one day enable the activation or repression of individual target genes in tumor tissue. In addition, genes identified through analysis of methylation may be targeted through other treatment modalities such as small molecules, biologics, and nucleic acid-based medicines.

## Acknowledgments

The results published here are in whole or part based upon data generated by the TCGA Research Network: https://www.cancer.gov/tcga.

Thanks to Amit Kulkarni for suggestions on TCGA data, to Dr. William Forrest for his input on statistical analysis, and to Dr. Jehnna Ronan for early advice and suggestions for direction.

## Supplementary Information

**S1 Fig. Most genes’ expression is inversely correlated with their methylation.** Histograms of Spearman correlation between gene methylation and RNA-Seq expression values. Each panel shows results for one tumor type. Tumor type abbreviations are in Table 1. Tumor types with fewer than 5 gene associations were excluded.

**S1 Table. Genes for which methylation is significantly associated with shorter survival within each study and tumor type (*q* < 0.05).** Each column contains the genes identified in one tumor type. The top row identifies the tumor types. Tumor type abbreviations are shown in Table 1.

**S2 Table. Genes for which methylation is significantly associated with greater survival within each study and tumor type (*q* < 0.05).** Each column contains the genes identified in one tumor type. The top row identifies the tumor types. Tumor type abbreviations are shown in Table 1.

**S3 Table. Genes in each of the four peaks of highest chromosomal density shown in figures 2 and 3.** Each column contains the genes under the peak on the chromosome specified in the first row.

**S4 Table. Overlap with Gene Ontology gene sets for genes associated with shorter survival across three or more studies.** Columns represent gene sets and rows represent genes. An X indicates that a gene was part of a gene set.

**S5 Table. Overlap with Gene Ontology gene sets for genes associated with greater survival across three or more studies.** Columns represent gene sets and rows represent genes. An X indicates that a gene was part of a gene set.

