## Supplementary Figure 1 for "Pan-Cancer Analysis of DNA Methylation Identifies Genes and Biological Functions Associated with Overall Survival"

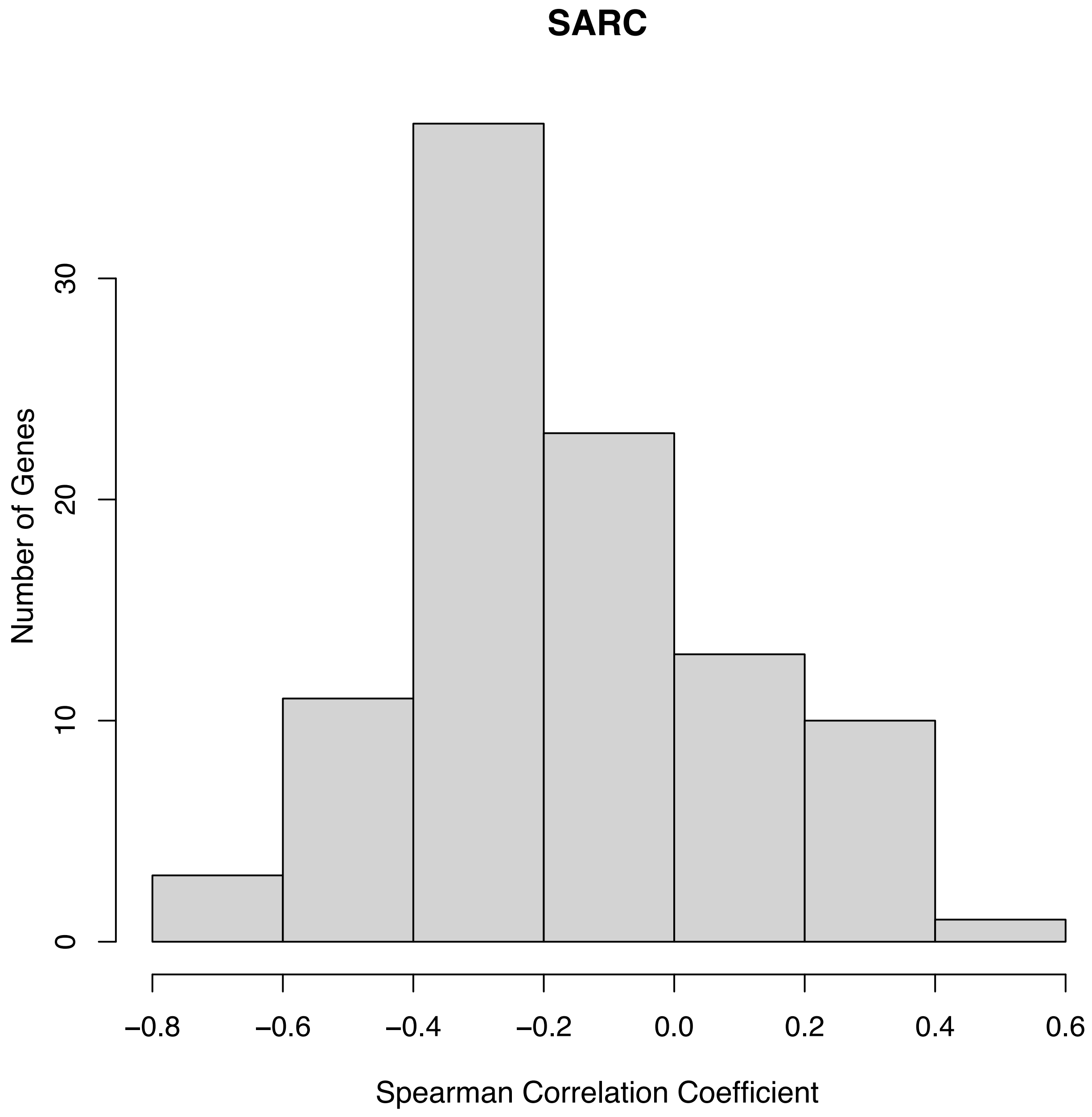
SARC
Number of Genes
0
10
20
30
−0.8
−0.6
−0.4
−0.2
0.0
0.2
0.4
0.6
Spearman Correlation Coefficient

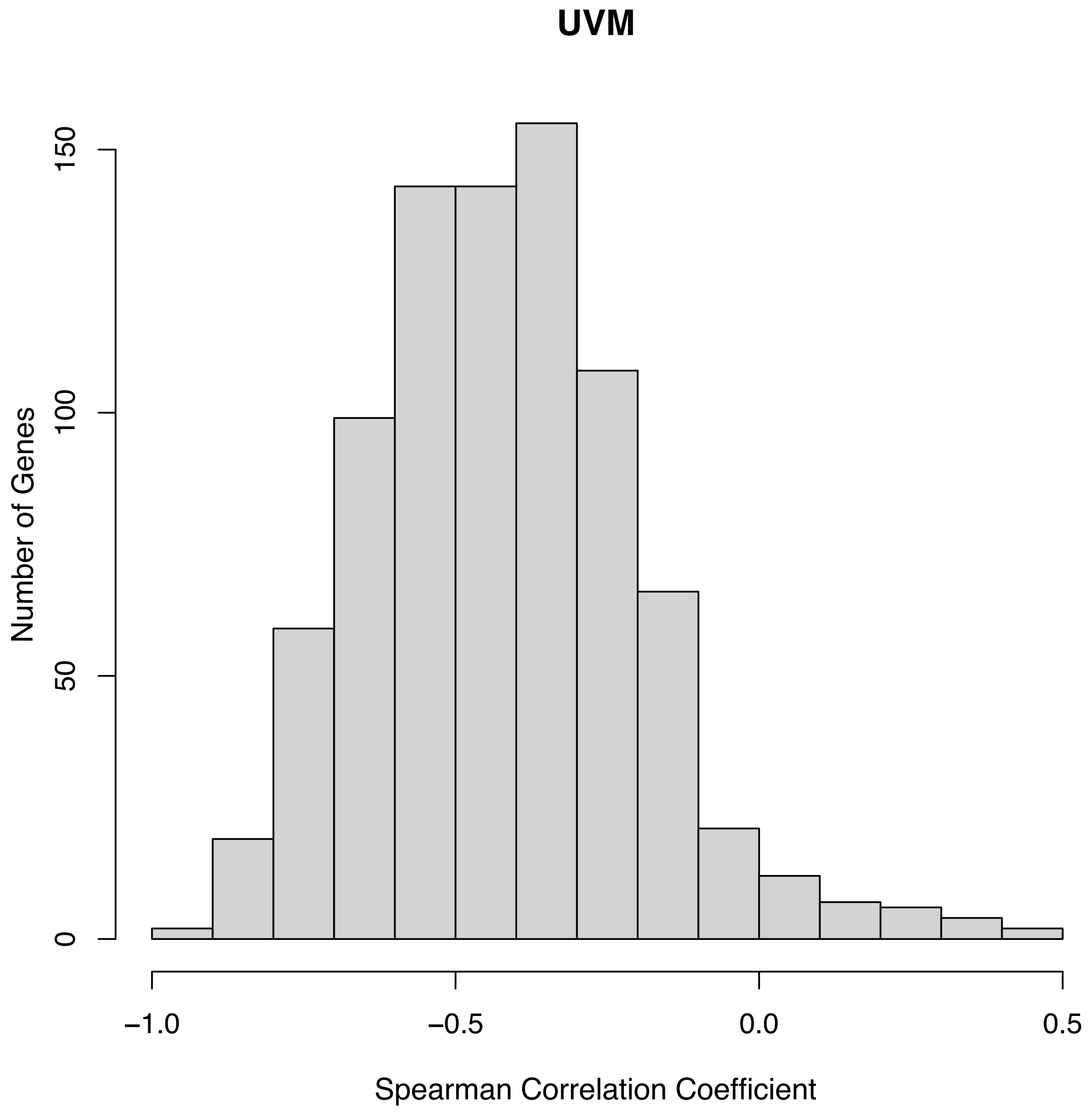
UVM
Number of Genes
0
50
100
150
−1.0
−0.5
0.0
0.5
Spearman Correlation Coefficient

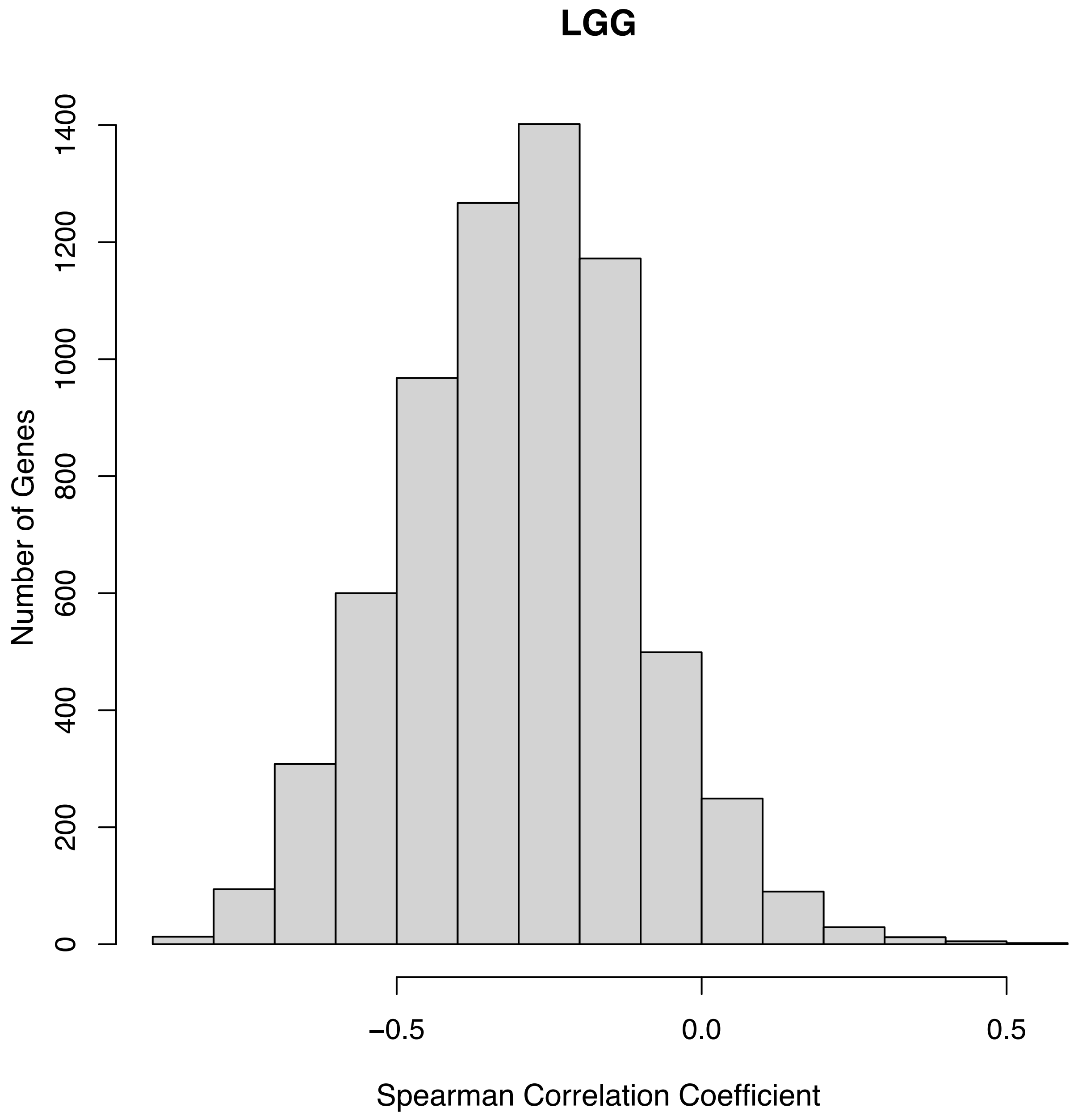

LGG
Number of Genes
0
200
400
600
800
1000
1200
1400
−0.5
0.0
0.5
Spearman Correlation Coefficient

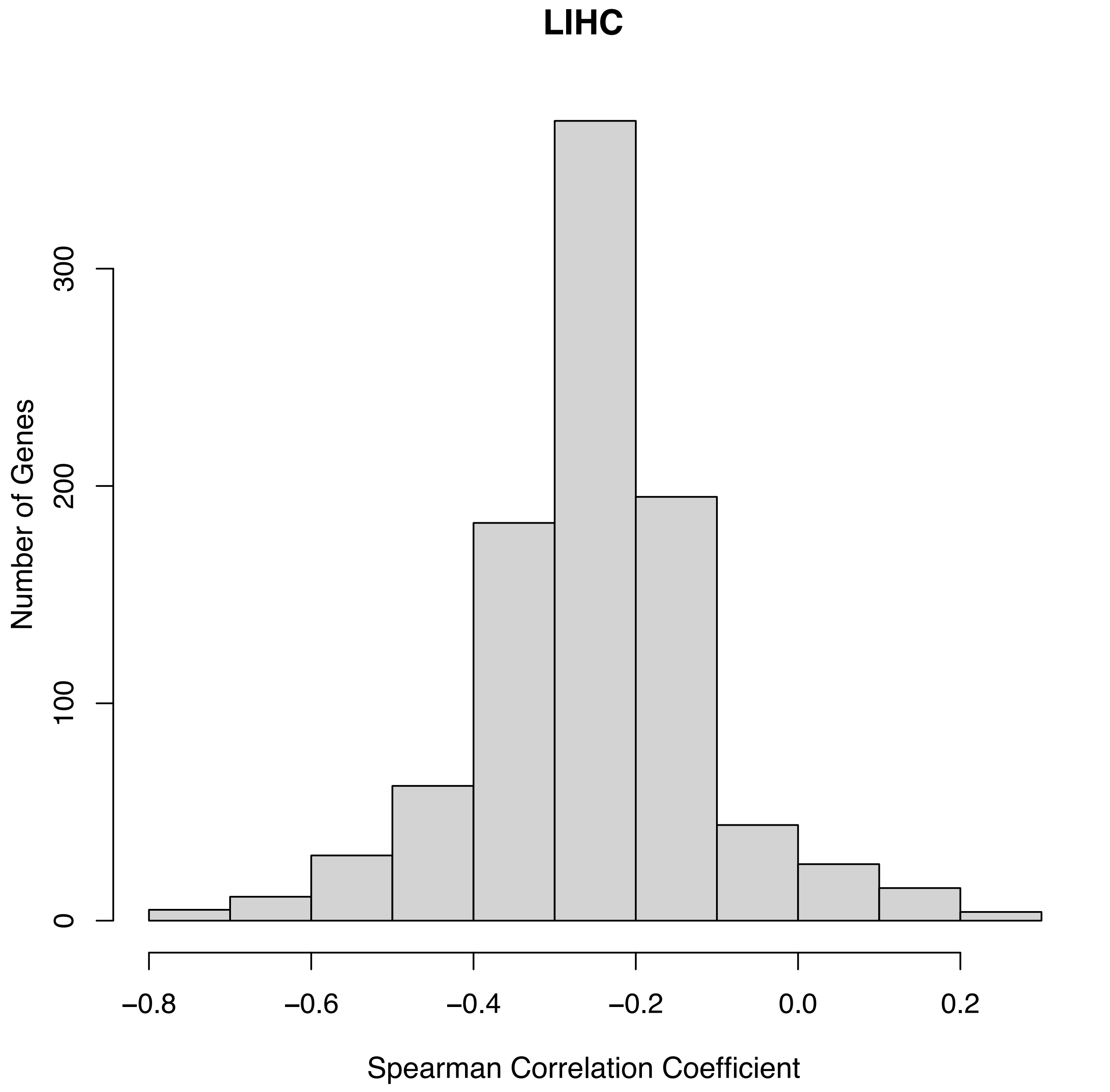
LIHC
Number of Genes
0
100
200
300
−0.8
−0.6
−0.4
−0.2
0.0
0.2
Spearman Correlation Coefficient

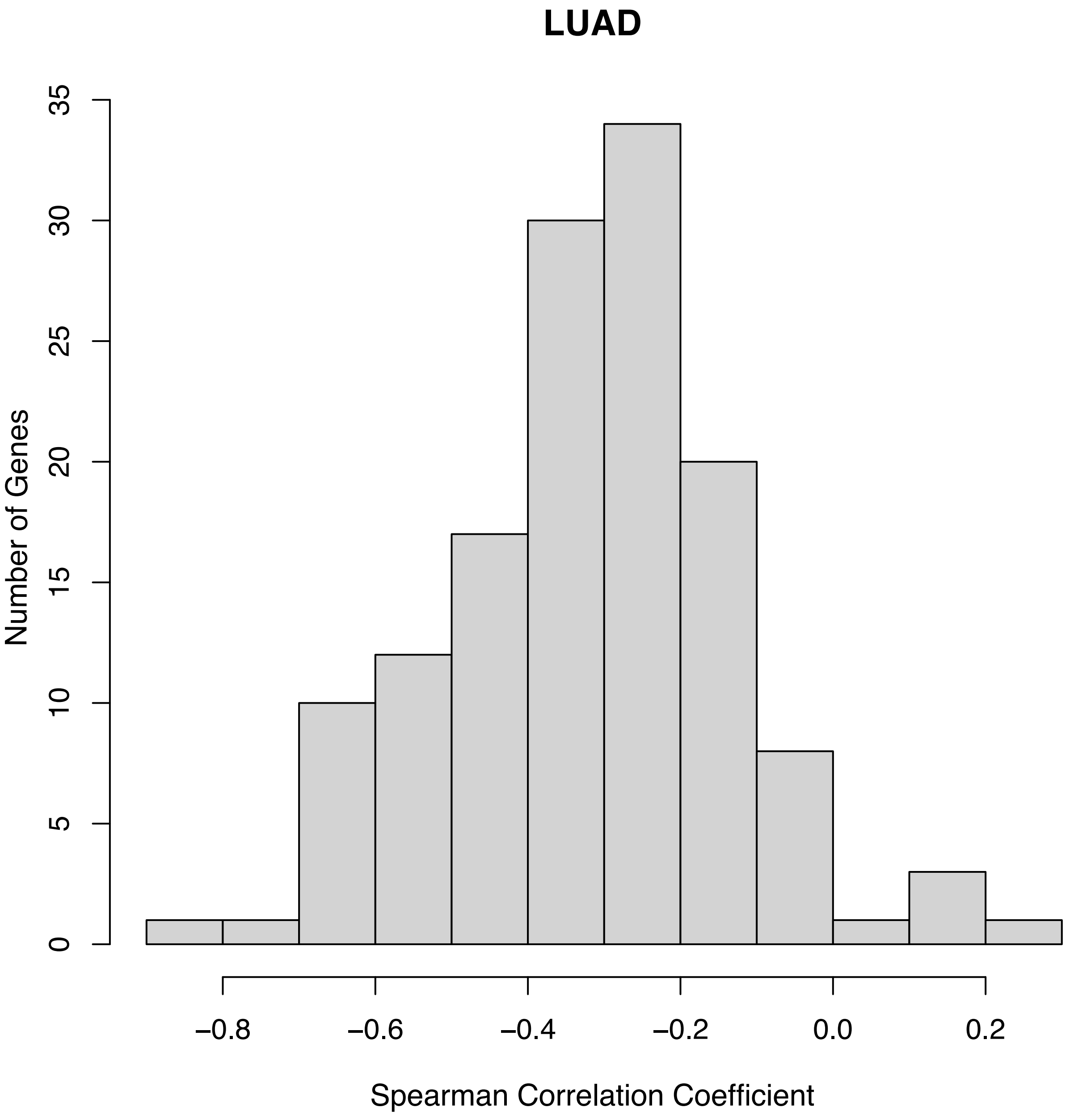
LUAD
Number of Genes
0
5
10
15
20
25
30
35
−0.8
−0.6
−0.4
−0.2
0.0
0.2
Spearman Correlation Coefficient

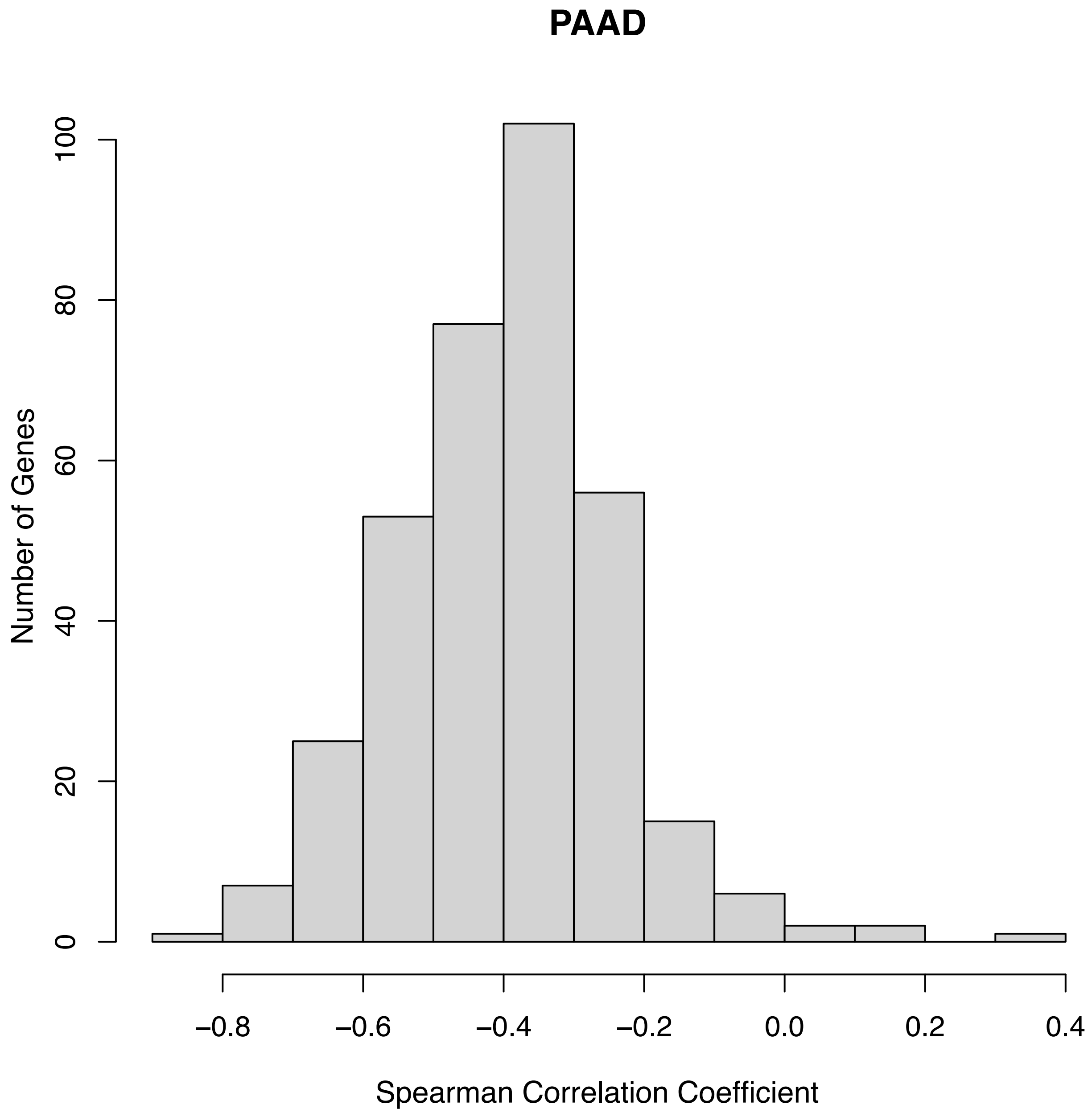

PAAD
Number of Genes
0
20
40
60
80
100
−0.8
−0.6
−0.4
−0.2
0.0
0.2
0.4
Spearman Correlation Coefficient

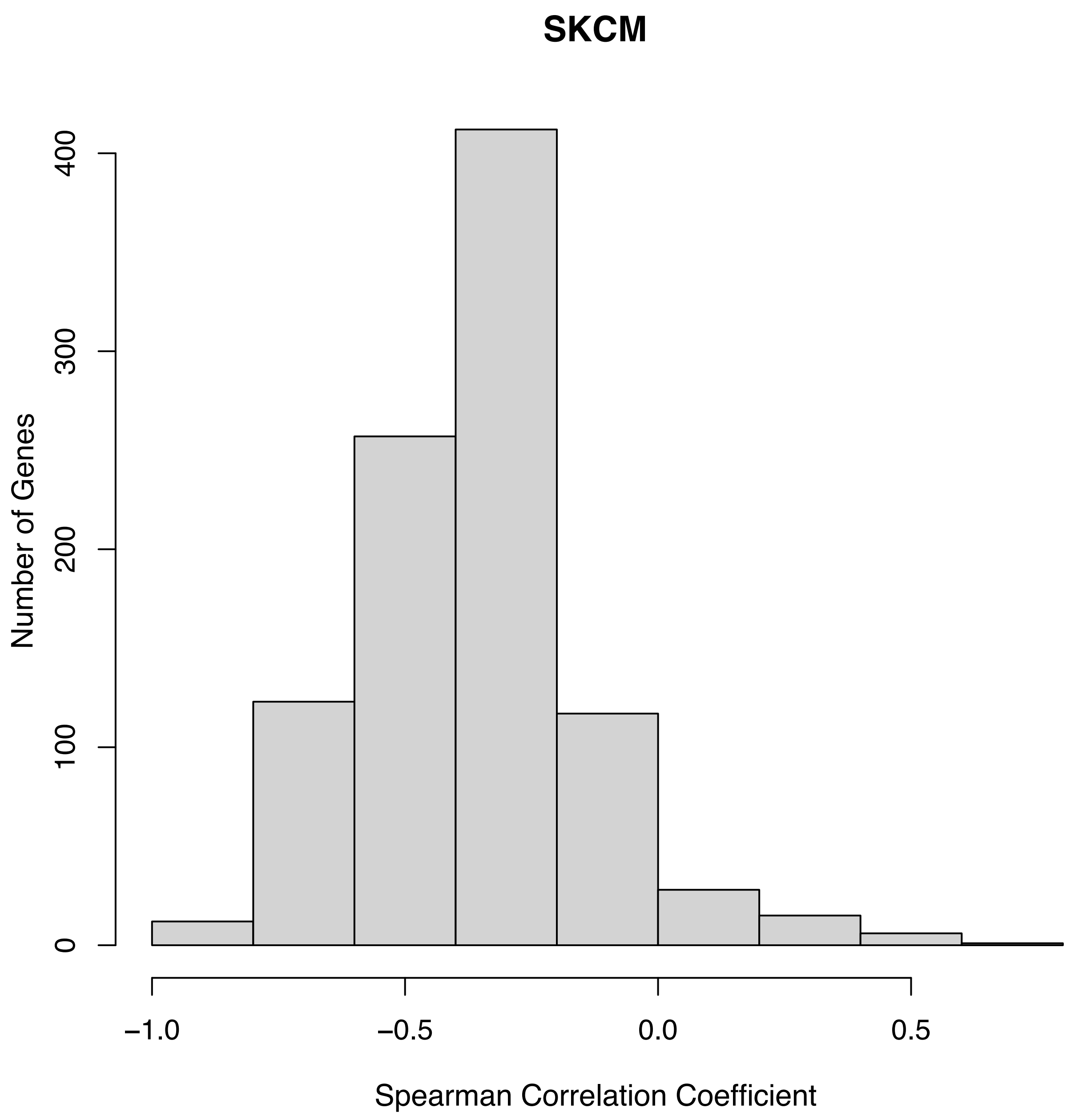
SKCM
Number of Genes
0
100
200
300
400
−1.0
−0.5
0.0
0.5
Spearman Correlation Coefficient

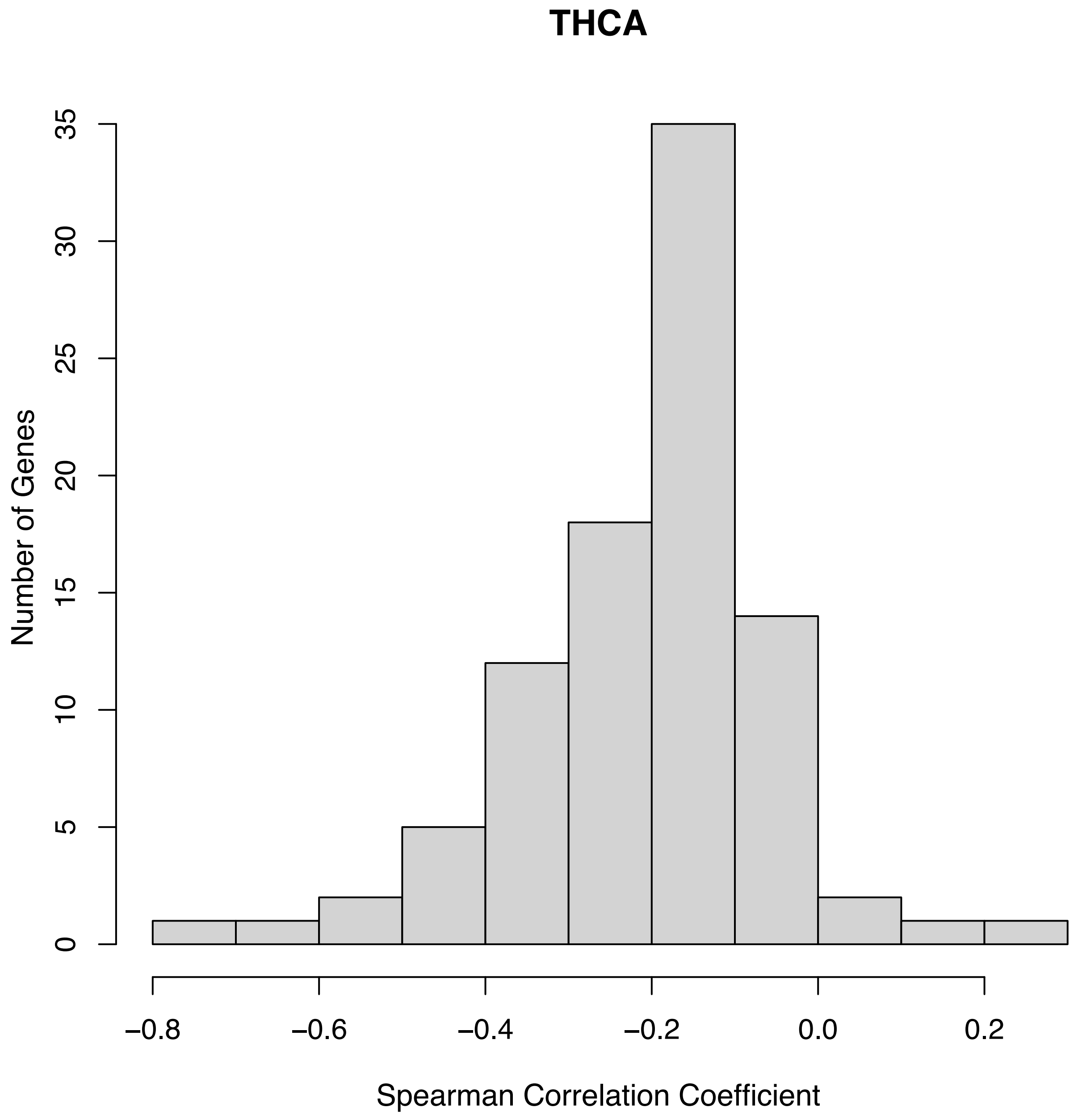

THCA
Number of Genes
0
5
10
15
20
25
30
35
−0.8
−0.6
−0.4
−0.2
0.0
0.2
Spearman Correlation Coefficient

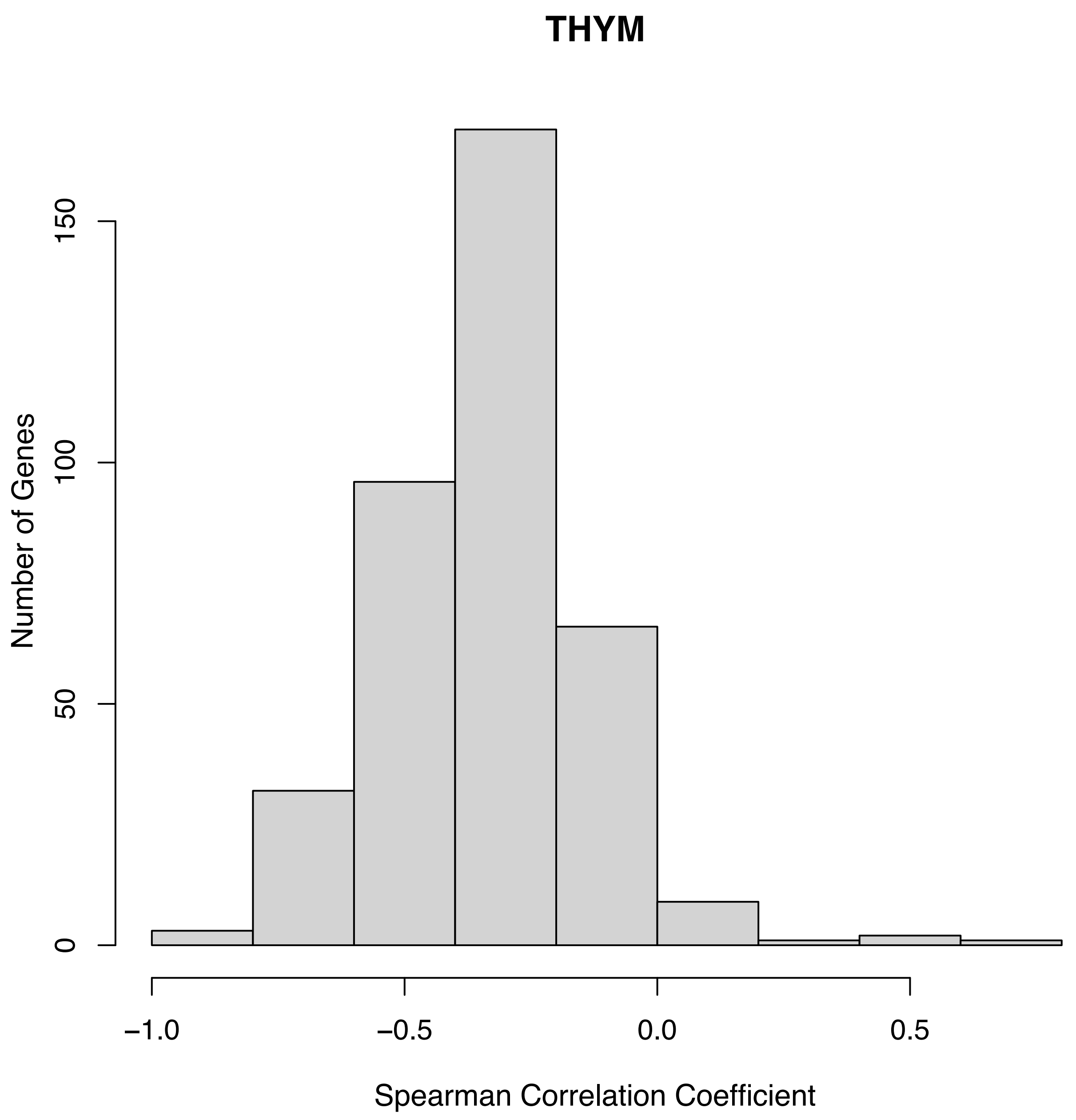

THYM
Number of Genes
0
50
100
150
−1.0
−0.5
0.0
0.5
Spearman Correlation Coefficient

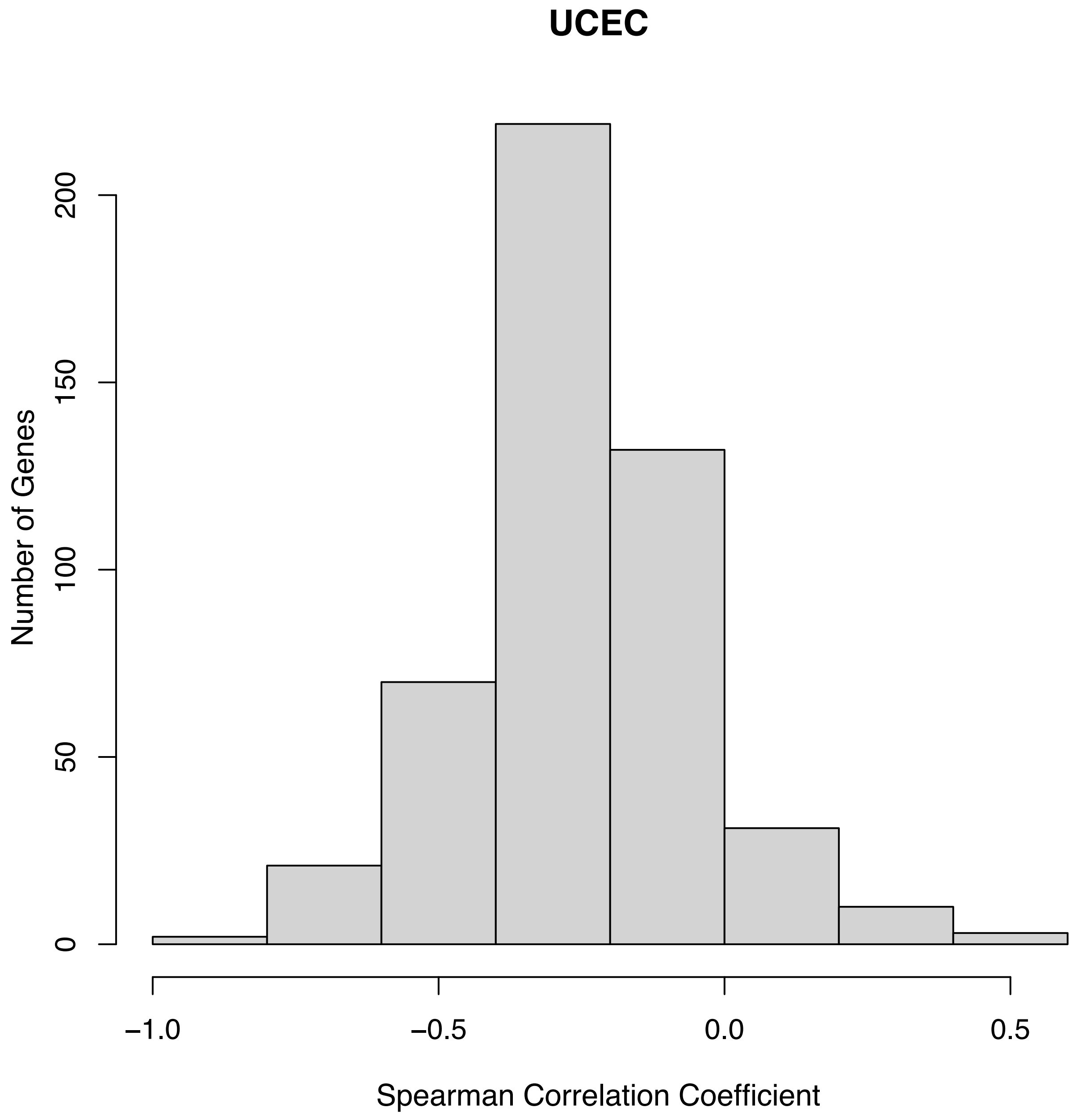
UCEC
Number of Genes
0
50
100
150
200
−1.0
−0.5
0.0
0.5
Spearman Correlation Coefficient

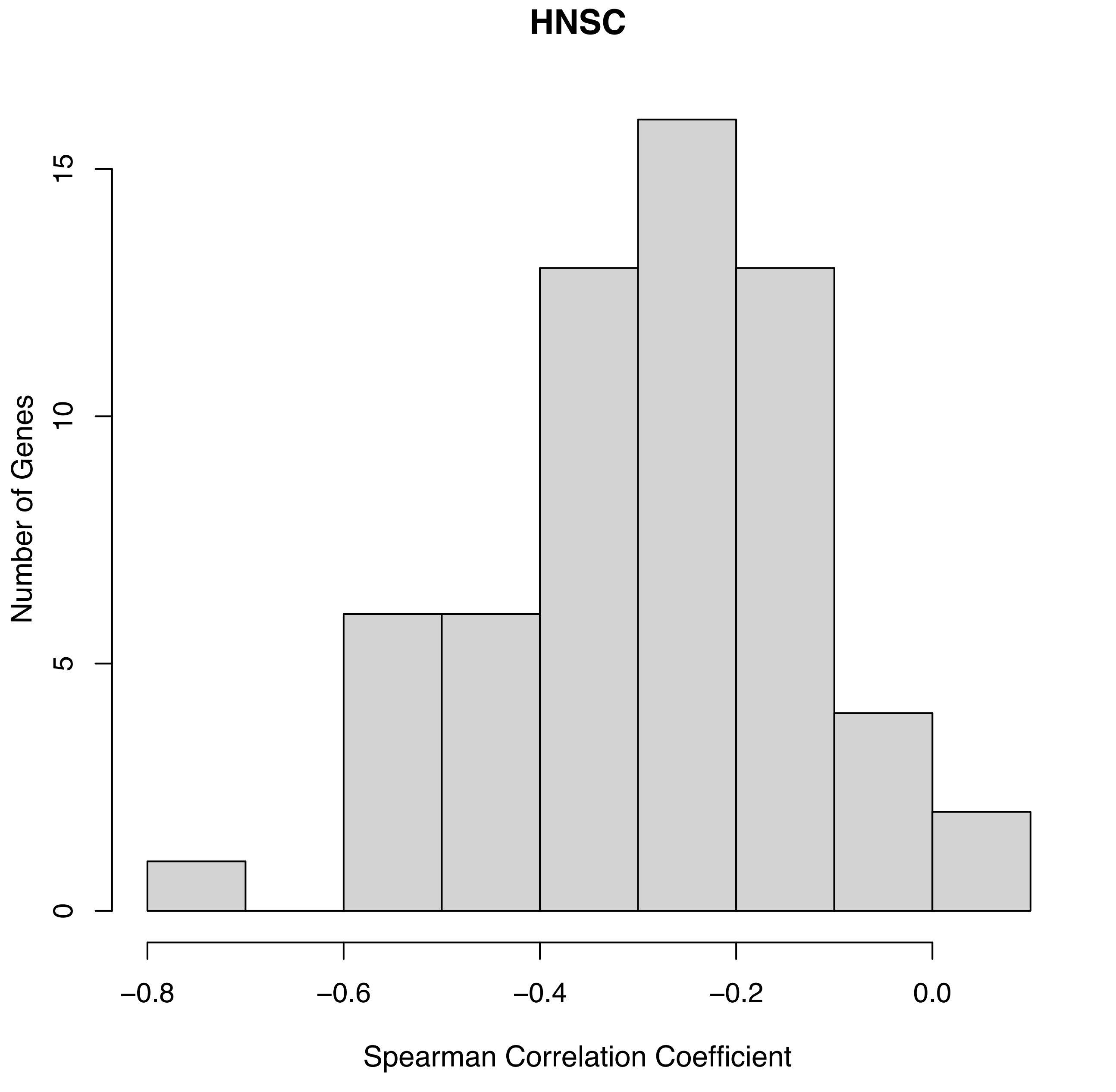
HNSC
Number of Genes
0
5
10
15
−0.8
−0.6
−0.4
−0.2
0.0
Spearman Correlation Coefficient

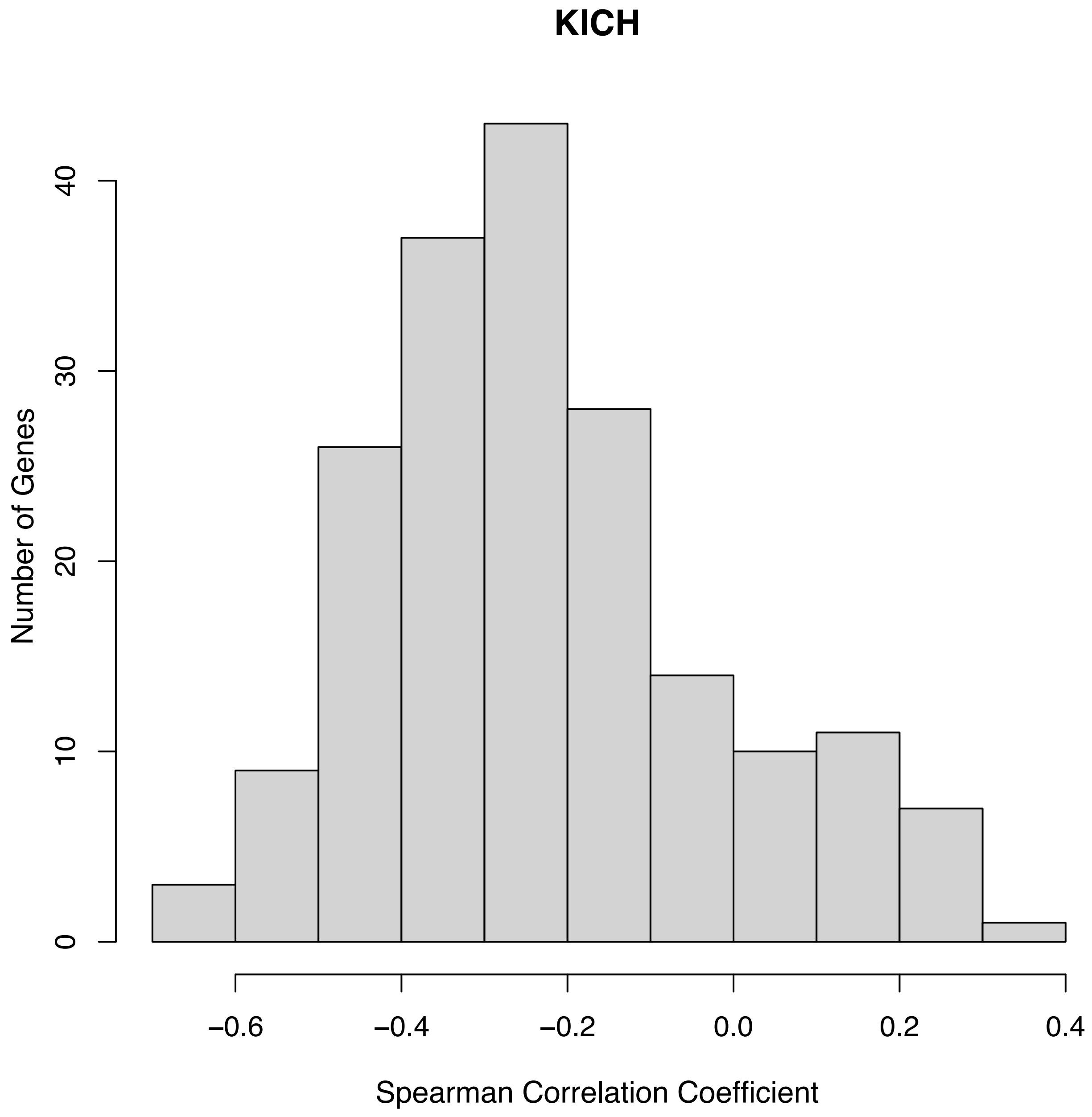
KICH
Number of Genes
0
10
20
30
40
−0.6
−0.4
−0.2
0.0
0.2
0.4
Spearman Correlation Coefficient

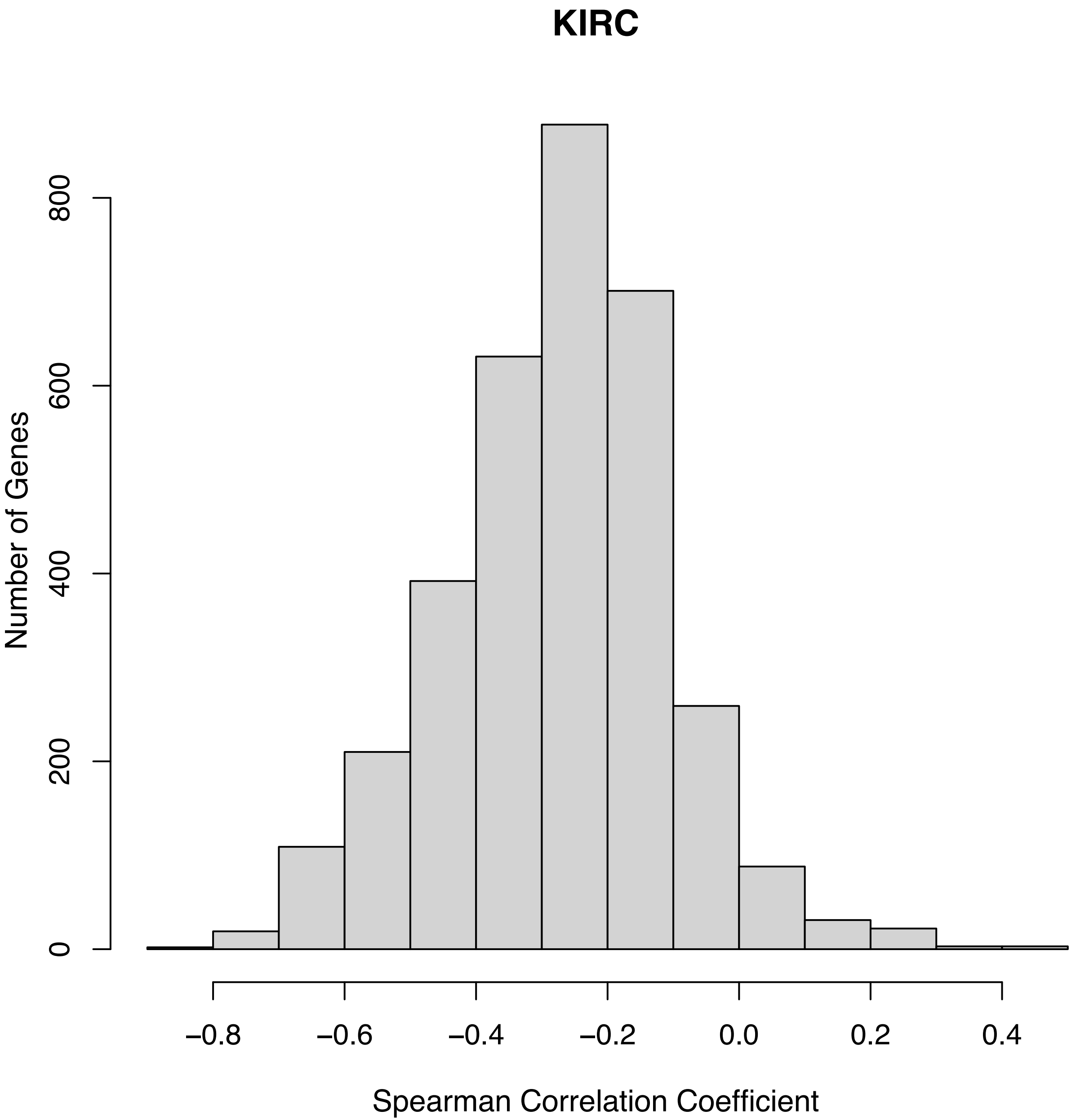

KIRC
Number of Genes
0
200
400
600
800
−0.8
−0.6
−0.4
−0.2
0.0
0.2
0.4
Spearman Correlation Coefficient

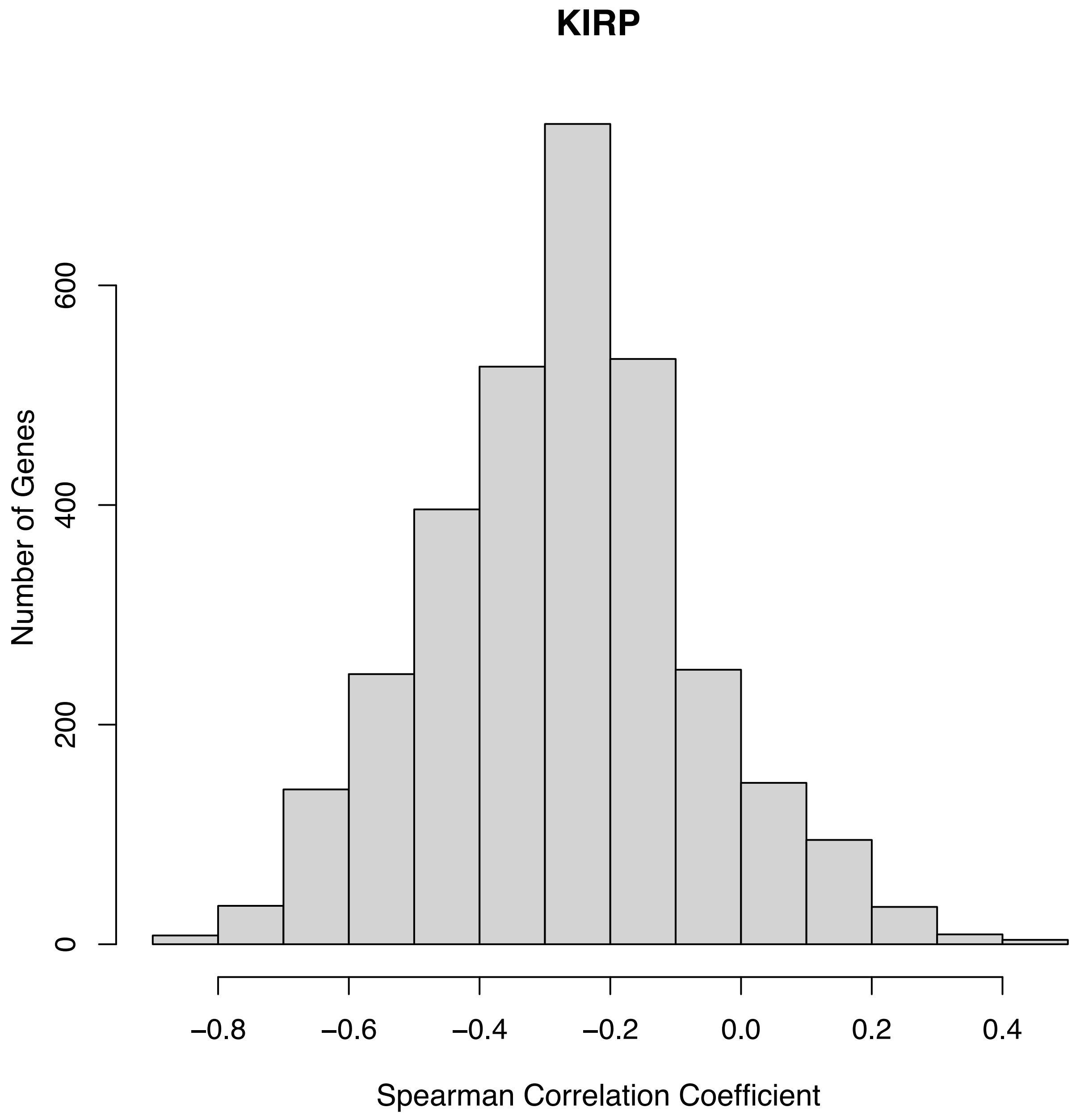

KIRP
Number of Genes
0
200
400
600
−0.8
−0.6
−0.4
−0.2
0.0
0.2
0.4
Spearman Correlation Coefficient

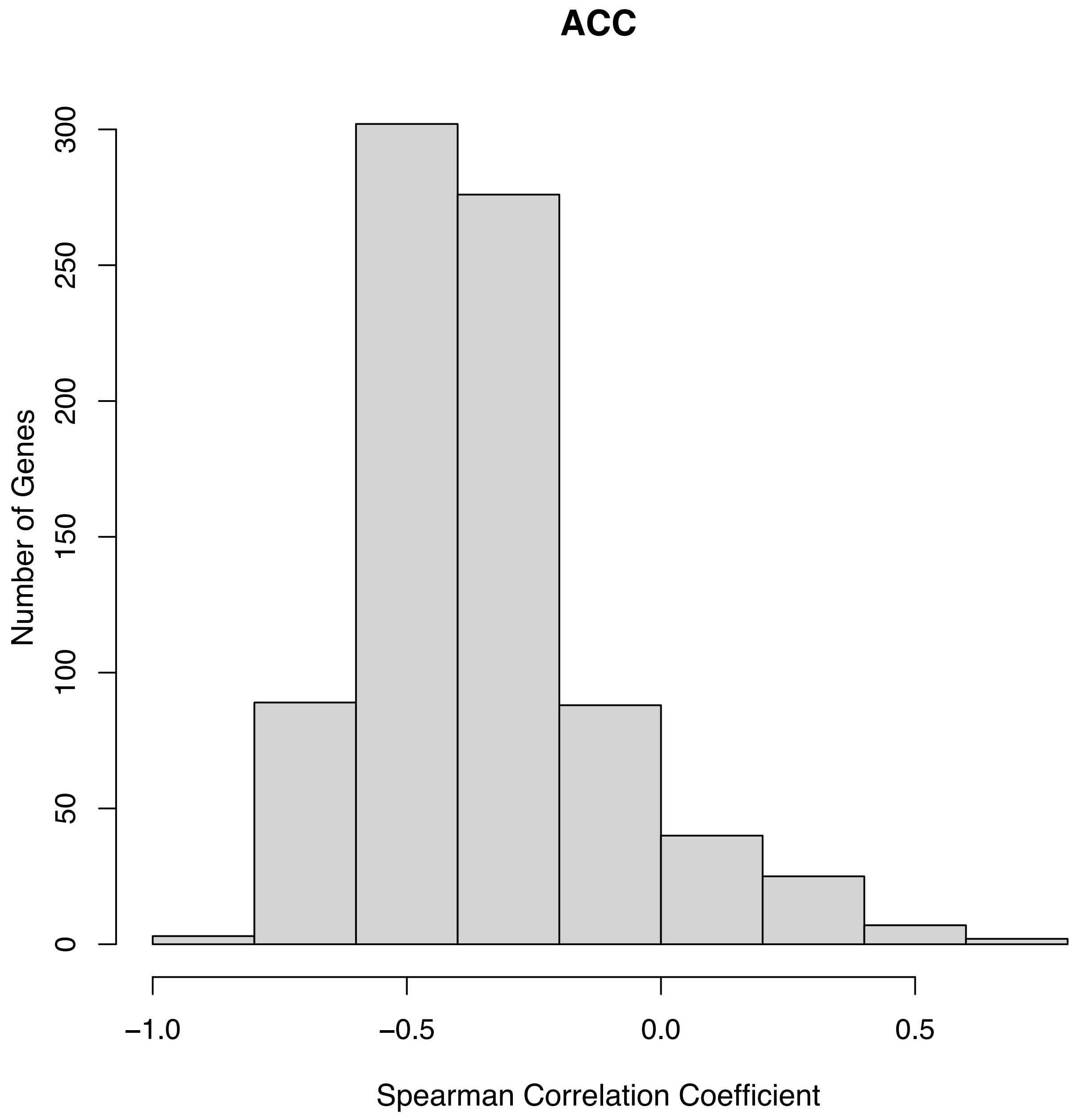

ACC
Number of Genes
0
50
100
150
200
250
300
−1.0
−0.5
0.0
0.5
Spearman Correlation Coefficient

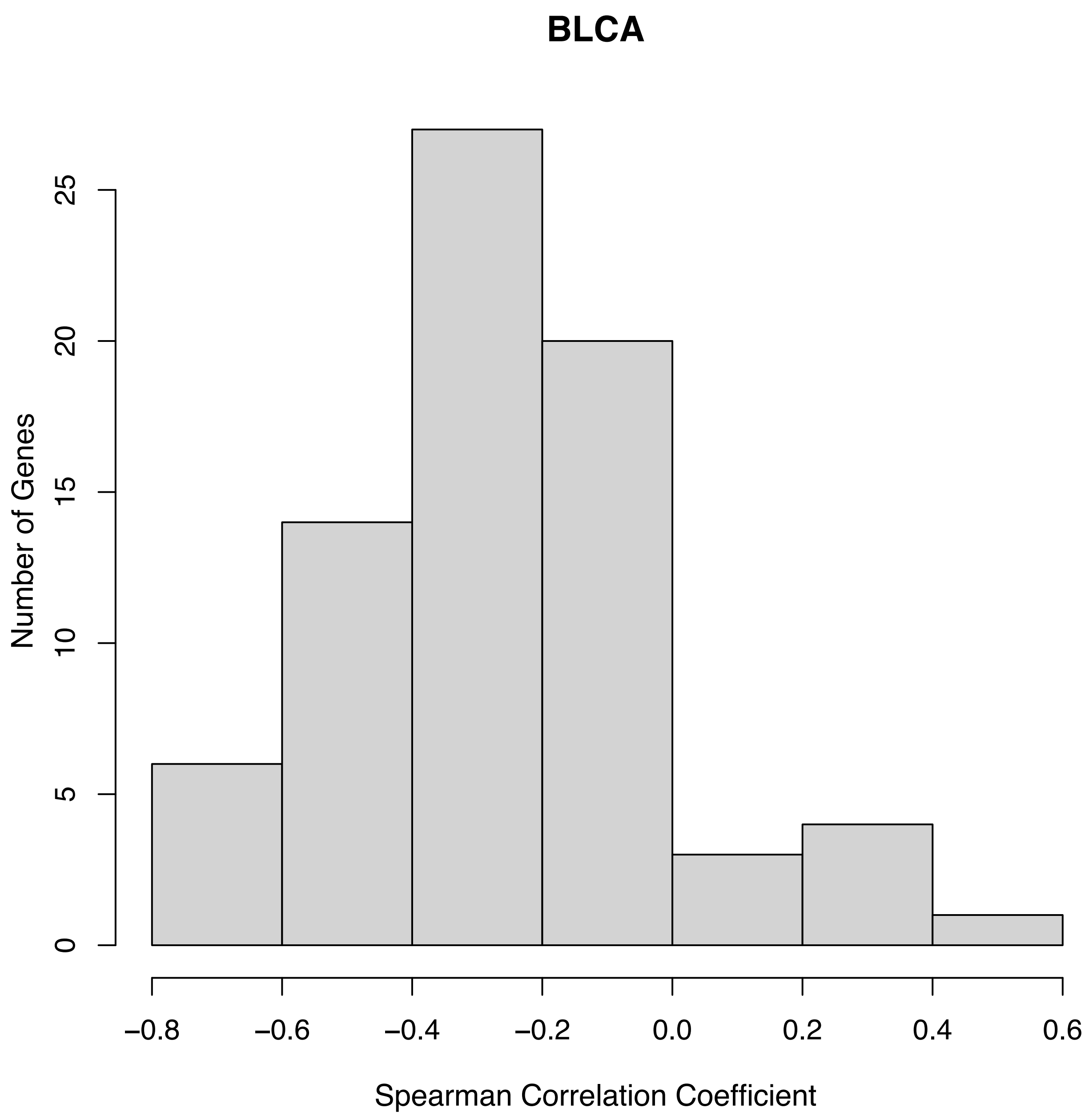
BLCA
Number of Genes
0
5
10
15
20
25
−0.8
−0.6
−0.4
−0.2
0.0
0.2
0.4
0.6
Spearman Correlation Coefficient

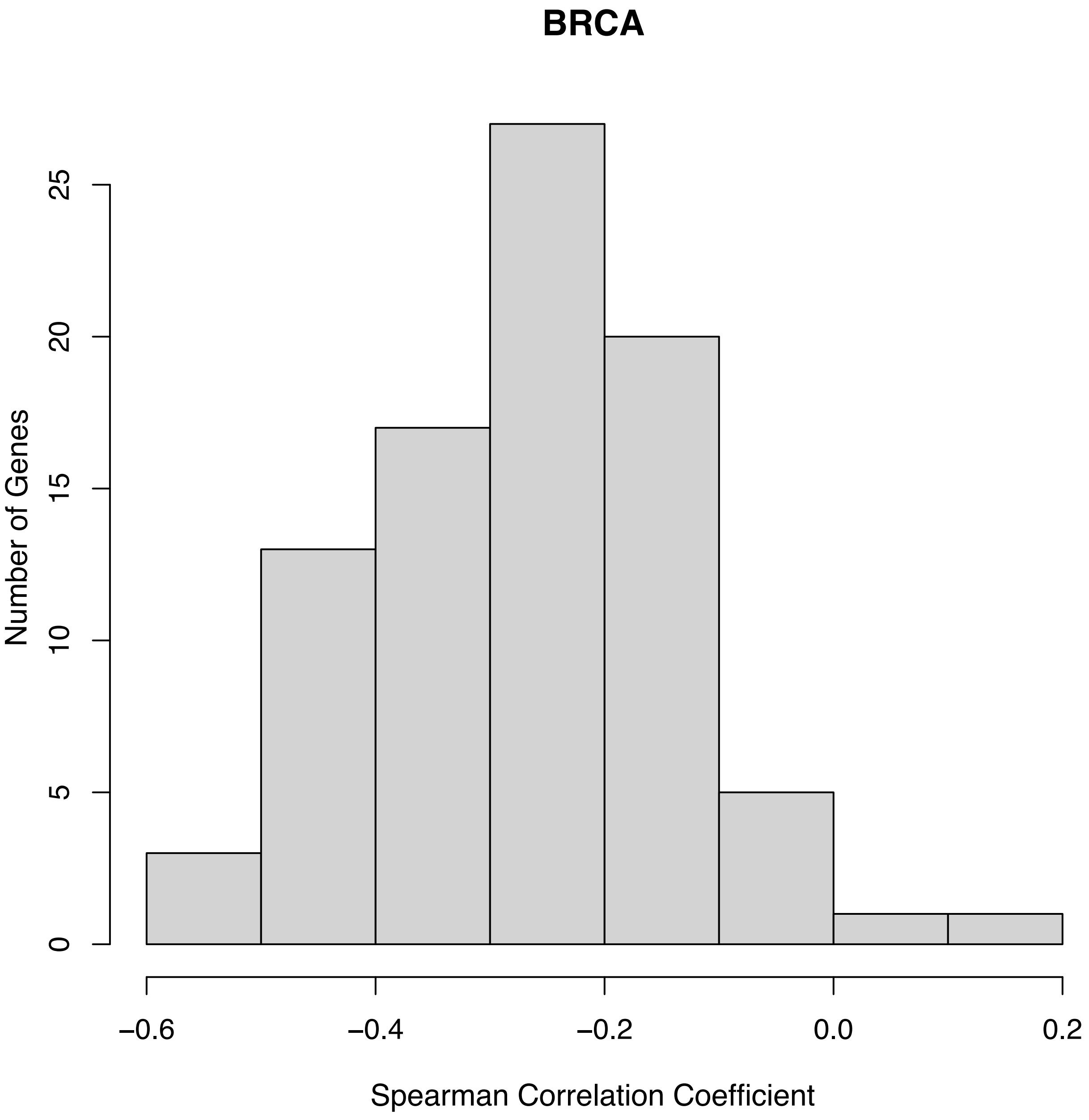
BRCA
Number of Genes
0
5
10
15
20
25
−0.6
−0.4
−0.2
0.0
0.2
Spearman Correlation Coefficient

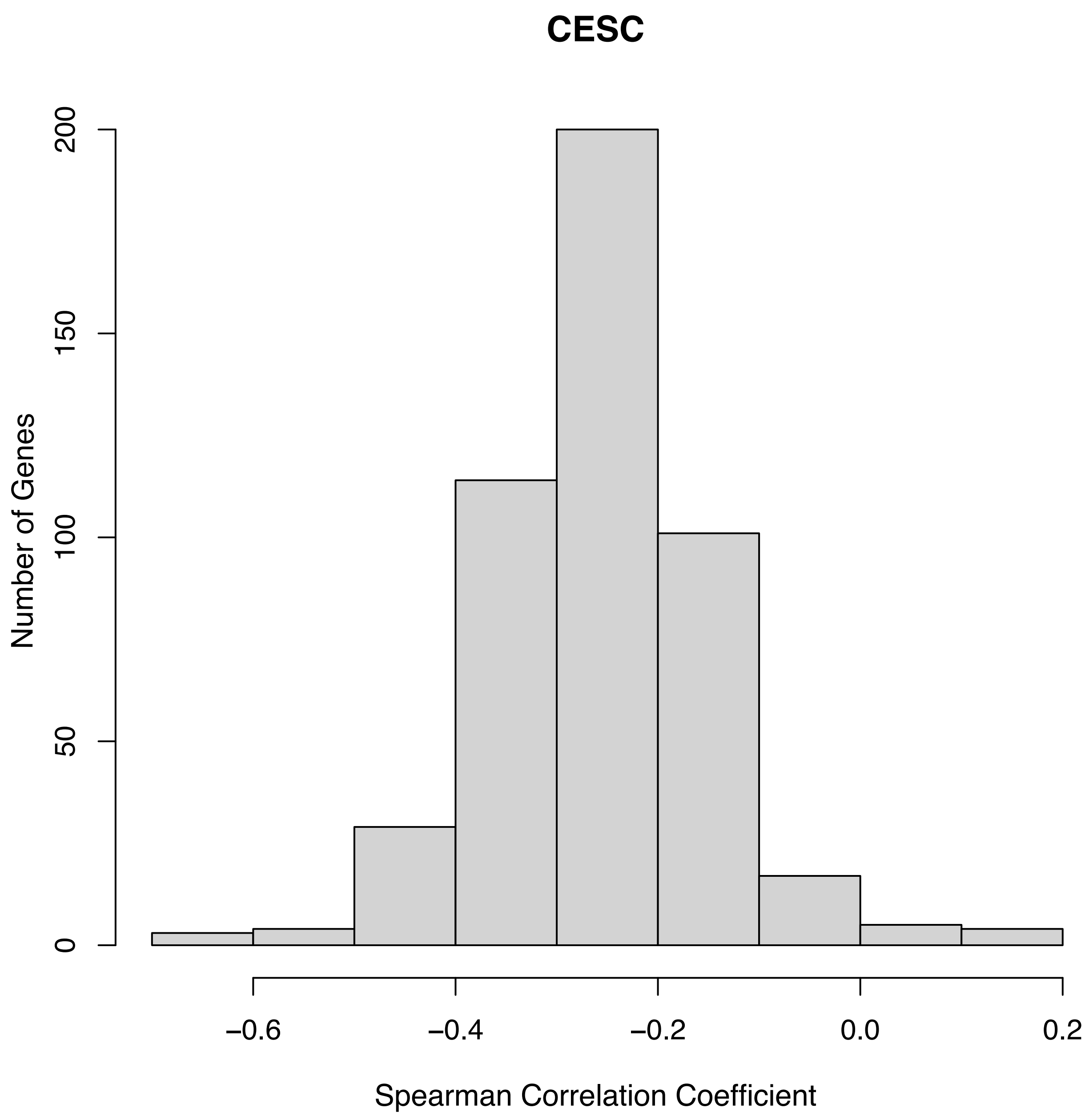
CESC
Number of Genes
0
50
100
150
200
−0.6
−0.4
−0.2
0.0
0.2
Spearman Correlation Coefficient
